# Common and distinct neural mechanisms of aversive and appetitive pain-related learning

**DOI:** 10.1101/2025.04.02.646781

**Authors:** Jialin Li, Katharina Schmidt, Lea Busch, Katarina Forkmann, Tamas Spisak, Jaspreet Kaur, Frederik Schlitt-Nguyen, Katja Wiech, Ulrike Bingel

## Abstract

Appetitive and aversive conditioning are both fundamental to adaptive behaviour, yet there remains limited understanding of how they differ on the behavioural and neural level. We investigated the two processes during acquisition and extinction using functional magnetic resonance imaging and behavioural measures. In a within-subject differential conditioning paradigm (preregistration DRKS00027448), aversive learning was induced by pairing visual cues with a temperature increase (pain rise), while appetitive learning involved a temperature decrease (pain reduction).

Valence and contingency ratings confirmed successful learning for both types of learning, though only the appetitive condition showed a return to baseline ratings during extinction, suggesting incomplete extinction in the aversive condition. On the neural level, both engaged the visual cortex during acquisition (with increased functional connectivity with the right frontal operculum) and the ventromedial prefrontal cortex (vmPFC) during extinction. However, aversive learning showed a stronger activation increase in the mediodorsal thalamus with heightened connectivity with the locus coeruleus during acquisition, as well as sustained parahippocampal activity during extinction. Moreover, incomplete extinction in the aversive condition (as indicated by contingency ratings) was associated with sustained activity in the visual cortex during pain anticipation.

These results suggest that while appetitive and aversive learning share activation in regions involved in sensory processing (occipital lobe) and learning (vmPFC), aversive learning uniquely engages areas promoting rapid acquisition (mediodorsal thalamus and locus coeruleus) and cautious unlearning, in line with the notion of a ‘better-safe-than-sorry’ strategy.

## Introduction

Human behaviour can be seen as guided by two key principles: avoiding harm and seeking what will ensure survival. Thus, learning from cues that signal potential harm (aversive learning) is as crucial as learning from cues that lead to desirable outcomes (appetitive learning). Both types of learning have been extensively studied using conditioning paradigms, where aversive or rewarding stimuli (unconditioned stimuli, US) are repeatedly paired with predictive cues (conditioned stimuli, CS) in the acquisition phase. Previous research has implicated different brain regions in these processes, with ‘fear circuits’ including brain regions such as dorsal anterior cingulate cortex (dACC) and anterior insula engaged in aversive learning^1–3^ and mesocorticolimbic reward circuits involved in appetitive learning^4–7^. Although some studies have explored common mechanisms, for instance by examining neural similarities between fear and reward conditioning across paradigms^8^, important communalities and differences may be overlooked when the two processes are studied separately and in different modalities (e.g., electric shock vs. monetary reward)^3,8^.

Pain offers an ecologically valid model for studying aversive and appetitive learning within the same paradigm. Due to its aversive nature, pain naturally captures attention and drives learning to avoid future harm^9,10^. Conversely, the alleviation of pain can be regarded a rewarding experience, governed by the principles of appetitive learning^11^. In a seminal study, Seymour et al. (2005) examined both pain and relief learning in the same sample and revealed that expectations of pain and relief engage opposing brain networks^12^. While this study provided valuable insights into the acquisition of cue-outcome associations, it did not address another key aspect of learning – the modification of learned associations when new information becomes available. When the cue is no longer followed by pain or relief, the CS-US associations must be updated to allow for the meaningful prediction of future events. Difficulties in both acquisition and extinction learning are central to a range of disorders including anxiety disorders and chronic pain and have important implications for treatment efficacy^11,13^. In a recent behavioural study in healthy individuals, we used an established capsaicin-induced tonic heat pain model that pairs predictive cues with either pain enhancement or reduction to investigate both aversive and appetitive pain-related learning within the same paradigm and sample. The results demonstrated stronger acquisition during aversive learning and incomplete extinction for aversive events^14^, suggestive of a “better-safe-than-sorry” strategy to minimise harm.

Here, we employed the same paradigm to characterise the joint and distinct neural underpinnings of both types of learning during acquisition and extinction in healthy individuals. Thermal stimulation was applied to a capsaicin-pretreated area on the forearm. Temperatures were individually calibrated to maintain moderate tonic pain throughout the experiment, with pain increase and pain decrease to induce pain exacerbation and relief, respectively. During the acquisition phase, geometrical cues (CS_increase_, CS_decrease_, CS_medium_) signalled pain exacerbation (US_increase_), pain decrease (US_decrease_) and no change in temperature (US_medium_), respectively, whereas all cues were followed by US_medium_ during the extinction phase (Fig. 1). CS-US learning was assessed through valence ratings to evaluate the affective dimension and contingency ratings to measure conscious learning. Participants also provided US pain intensity and (un)pleasantness ratings. Brain activity was recorded using functional magnetic resonance imaging (fMRI) during both the acquisition and extinction phase. We hypothesised that (1) CS-US associations for aversive events are acquired faster and extinguish slower than for appetitive events (i.e., in steeper/flatter learning slopes); and (2) during both acquisition and extinction, aversive and appetitive learning engage shared and distinct neural networks.

**Figure 1.**
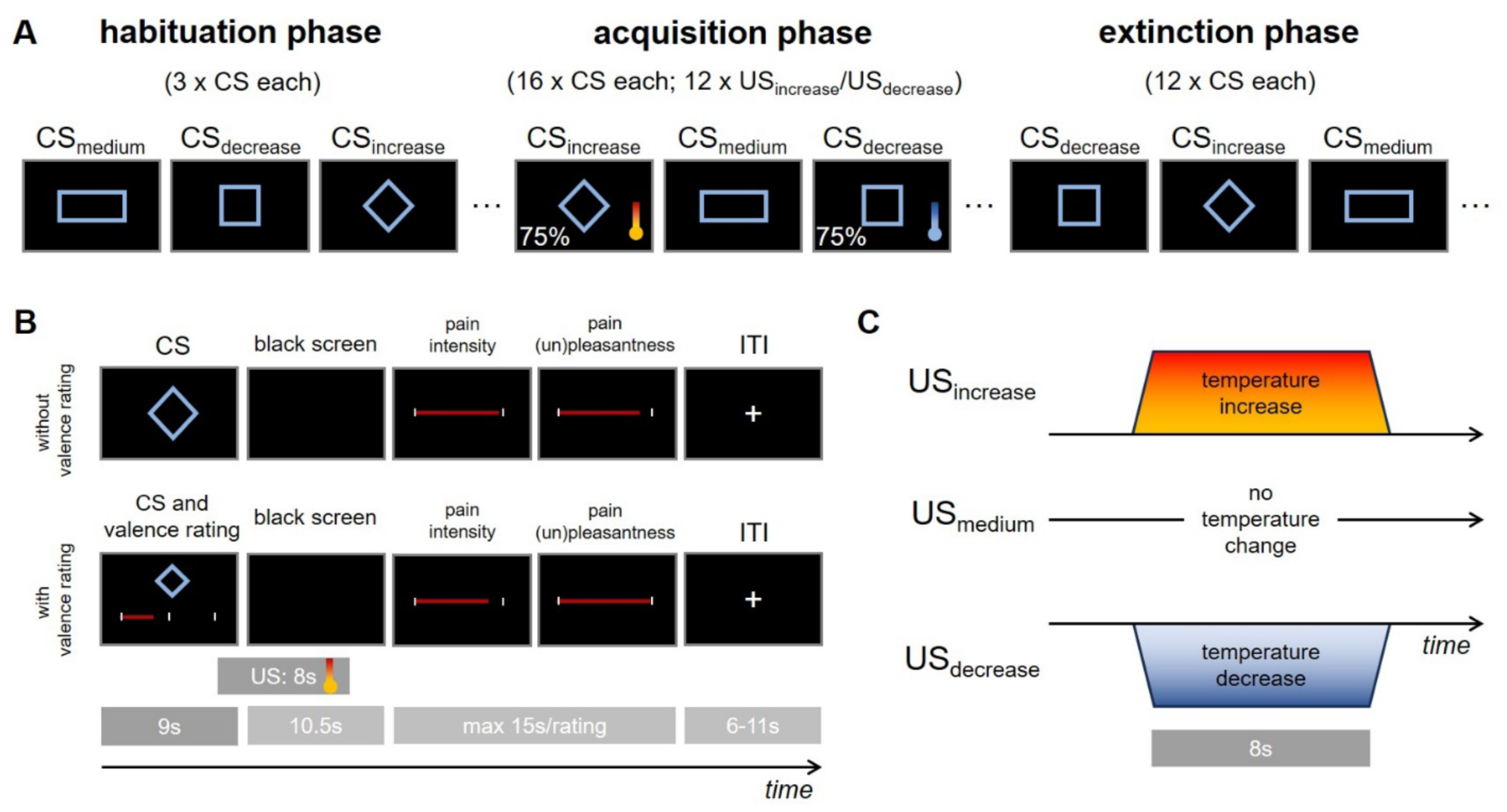
Study and trial design. (A) Study design with habituation, acquisition and extinction phase. During the acquisition phase, 75% of CSincrease and CSdecrease trials were followed by a temperature increase or temperature decrease, respectively. In the habituation and extinction phases, none of the CSs was reinforced. (B) Design of trials without (upper row) and with valence ratings (lower row) including pain intensity and (un)pleasantness ratings. Contingency ratings (not shown) were acquired at the end of the acquisition and extinction phase. (C) Temperature manipulation. The temperature of the thermal device was increased in the aversive condition (USincrease), decreased in the appetitive condition (USdecrease) and kept constant in the control condition (USmedium). CS = conditioned stimulus; US = unconditioned stimulus; ITI = inter-trial interval.

## Results

A total of *n* = 84 healthy individuals were enrolled in the study to participate in a differential conditioning task during functional magnetic resonance imaging (fMRI) to compare aversive and appetitive acquisition and extinction learning using an experimental pain model. Data from *n* = 16 participants were excluded from the behavioural analyses for several reasons: the thermal stimuli (US_increase_, US_decrease_ and US_medium_) could not be calibrated as intended or changed substantially during acquisition training (e.g., participants rated the US_medium_ as more painful than the US_increase_, *n* = 9), early termination of the experiment by the participants (*n* = 3) and technical errors (*n* = 4). The behavioural analyses are thus based on *n* = 68 participants (*n*_male_ = 33, *n*_female_ = 35, *Mean* age = 37.6, *SD* = 13.2). Further sample characteristics can be found in Supplementary Table S1. Of these 68 participants, *n* = 7 participants were excluded from the fMRI analysis due to missing data in either the brain structural or functional brain MRI images (*n* = 6) and the logfile (*n* = 1), resulting in a final imaging sample size of *n* = 61 (*n*_male_ = 31, *n*_female_ = 30, *Mean* age = 39.5, *SD* = 14.6).

In a within-subject design, participants learned to associate visual cues with an aversive increase (CS_increase_), an appetitive decrease (CS_decrease_), or no change (CS_medium_) in noxious thermal input during acquisition training. In the subsequent extinction training, none of the cues was followed by changes in noxious input (see Methods for details on the paradigm). Behavioural measures of learning included CS valence ratings and CS-US contingency ratings and we identified joint and distinct changes in blood oxygenation level-dependent (BOLD) signals and functional connectivity of regions-of-interests (ROIs) across aversive and appetitive learning. To control for spontaneously occurring changes in outcome measures unrelated to temperature manipulations, all analyses compared behavioural and neural responses in the aversive and appetitive conditions against the control condition.

Analyses of pain intensity and (un)pleasantness ratings obtained during acquisition training confirmed successful calibration of the noxious stimuli. Compared to the control condition, the thermal stimulation in the aversive condition was on average perceived as significantly more painful (US_increase_ > US_medium_, *p*< .001; Fig. 2A) and unpleasant (US_increase_ > US_medium_, *p* < .001) and the stimulation in the appetitive condition as less painful (US_decrease_ < US_medium_, *p* < .001; Fig. 2A) and less unpleasant (US_decrease_ < US_medium_, *p*< .001). For details on pain intensity and pain (un)pleasantness ratings see Supplementary Information.

### Behavioural evidence for acquisition and extinction learning

#### Valence ratings: acquisition

Differential valence ratings, reflecting learning at the affective level relative to the control condition, significantly changed over the course of acquisition training in both conditions (linear mixed model, LMM; *time:* β = 4.55, Χ^2^(1) = 35.96, *p* < .001; Fig. 2B). Relative to the control condition, neither the slope nor the absolute level differed between the aversive and the appetitive condition (*CS type*: Χ^2^(1) = 0.43, *p* = .511; *CS type* x *time*: Χ^2^(1) = 1.66, *p* = .198; Fig. 2B). To explore whether this finding was due to any changes in the control condition, we also analysed valence ratings separately for each of the three conditions (Fig. 2C). This model revealed a significant interaction between *CS type* and *time* (Χ^2^(2) = 108.00, *p* < .001). While valence ratings increased for the CS_increase_ (β = 4.69, *t*(746) = 9.00, *p* < .001), they decreased for the CS_decrease_ (β = -3.09, *t*(755) = -5.70, *p* < .001). Pairwise comparisons showed that slopes for the reinforced CS differed significantly from the CS_medium_ (CS_increase_ > CS_medium_, Δβ = 3.12, *t*(749) = 4.19, *p* < .001; CS_decrease_ < CS_medium_, Δβ = 4.66, *t*(753) = 6.14, *p* < .001). Valence ratings also increased slightly for the CS_medium_ (β = 1.57, *t*(751) = 2.96, *p* = .010). These results indicate acquisition learning for both reinforced CS types, but also a small change in valence of the non-reinforced CS_medium_. An exploratory direct comparison of the CS_increase_ and CS_decrease_ showed that individual slopes differed between the CS_incease_ and CS_decrease_ (paired t-test on individual slopes with slopes for CS_decrease_ calculated as x (-1) for direct comparison, CS_incease_ > CS_decrease_, *t*(67) = -2.23, *p* = .029, *d* = 0.27). This finding confirms that changes in valence ratings were more pronounced in the aversive than the appetitive condition.

As shown in Fig. 2C, valence ratings were rapidly adjusted early in the acquisition phase. To investigate this initial learning further, we applied a linear model to compare changes in valence ratings from habituation to the first reinforced acquisition trial (A1-H0) by *CS type.* This exploratory analysis revealed a significant main effect of *CS type* (*F*(2,173) = 12.43, *p* < .001). Specifically, while the CS_increase_ already significantly differed from the CS_medium_ (*t*(173) = 4.65, *p* < .001), the CS_decrease_ did not (*t*(173) = 0.82, *p* = .694; Fig. 2D). This indicates that participants made stronger early adjustments to their ratings in the aversive than the appetitive condition.

#### Valence ratings: extinction

During extinction, differential valence ratings decreased across conditions (*time*: β = -3.59, Χ^2^(1) = 25.41, *p* < .001), but the decrease did not differ between *CS types* (*CS type* x *time*: Χ^2^(1) = 0.86, *p* = .353; Fig. 2B). To test whether this finding could be explained by a change in the control condition, we analysed the valence ratings of all three conditions separately (Fig. 2C). While both CS_increase_ and CS_decrease_ showed a change in valence ratings towards the expected direction (i.e., decreasing ratings in CS_increase_ and increasing ratings for CS_decrease_), valence ratings for the CS_medium_ did not change significantly (β = 0.09, *t*(574) = 0.18, *p* = .997). An exploratory direct comparison of the CSincrease and CSdecrease showed that individual slopes did not differ between CS types (paired t-test on individual slopes, slopes for CSdecrease value x (-1), *t*(67) = -1.14, *p* = .257, *d* = 0.14).

#### CS-US contingency ratings

As a complementary measure of associative conditioning that assesses the cognitive aspect of learning, we also collected contingency ratings (i.e., estimated frequency of CS-US-associations) at the end of acquisition and extinction training, respectively. Analogously to the valence ratings, we hypothesised that acquisition learning would be stronger, and extinction would be weaker in the aversive condition.

The results showed a significant interaction of *CS type* and *time* (Χ^2^(2) = 12.57, *p* = .002; Fig. 2E). After acquisition training, contingency ratings for the CS_increase_ and the CS_decrease_ were significantly increased in the expected direction relative to the CS_medium_ (CS_increase_ > CS_medium_, β = 36.6, *t*(335) = 6.37, *p* < .001; CS_decrease_ > CS_medium_, β = 21.2, *t*(335) = 3.69, *p* < .001). Contingency ratings for both CS_increase_ and CS_decrease_ decreased significantly from after acquisition to after extinction training (extinction - acquisition: CS_increase_, β = -33.7, *t*(335) = -5.86, *p* < .001; CS_decrease_, β = -37.7, *t*(335) = -6.57, *p* < .001), and at trend level for the CS_medium_ (β = -11.0, *t*(335) = -1.92, *p* = .056). While the change of the CS_increase_ and the CS_decrease_ differed significantly from the change of the CS_medium_ (change CS_increase_ > CS_medium_, β = -22.68, *t*(335) = -2.79, *p* = .017; change CS_decrease_ > CS_medium_, β = -26.71, *t*(335) = -3.29, *p* = .003), the difference between CS_increase_ and CS_decrease_ did not reach significance (β = 4.03, *t*(335) = 0.50, *p* = .945).

Contingency ratings were higher for the increase condition than for the decrease condition at both time points (after acquisition, β = 15.4, *t*(335) = 2.67, *p* = .021; after extinction, β = 19.4, *t*(335) = 3.38, *p* = .002). While after extinction training, the CS_increase_ still differed from the CS_medium_ (CS_increase_ > CS_medium_, β = 13.9, *t*(335) = 2.42, *p* = .043), the CS_decrease_ and CS_medium_ did no longer differ significantly anymore (β = -5.5, *t*(335) = -0.96, *p* = .604). Importantly, the CS_increase_ remained significantly different from zero after extinction training (*t*(380) = 5.15, *p* < .001), while the CS_medium_ (*t*(380) = 1.92, *p* = .159) and the CS_decrease_ did not (*t*(380) = 0.64, *p =* .893), indicating incomplete extinction for contingency ratings for the aversive condition.

**Figure 2.**
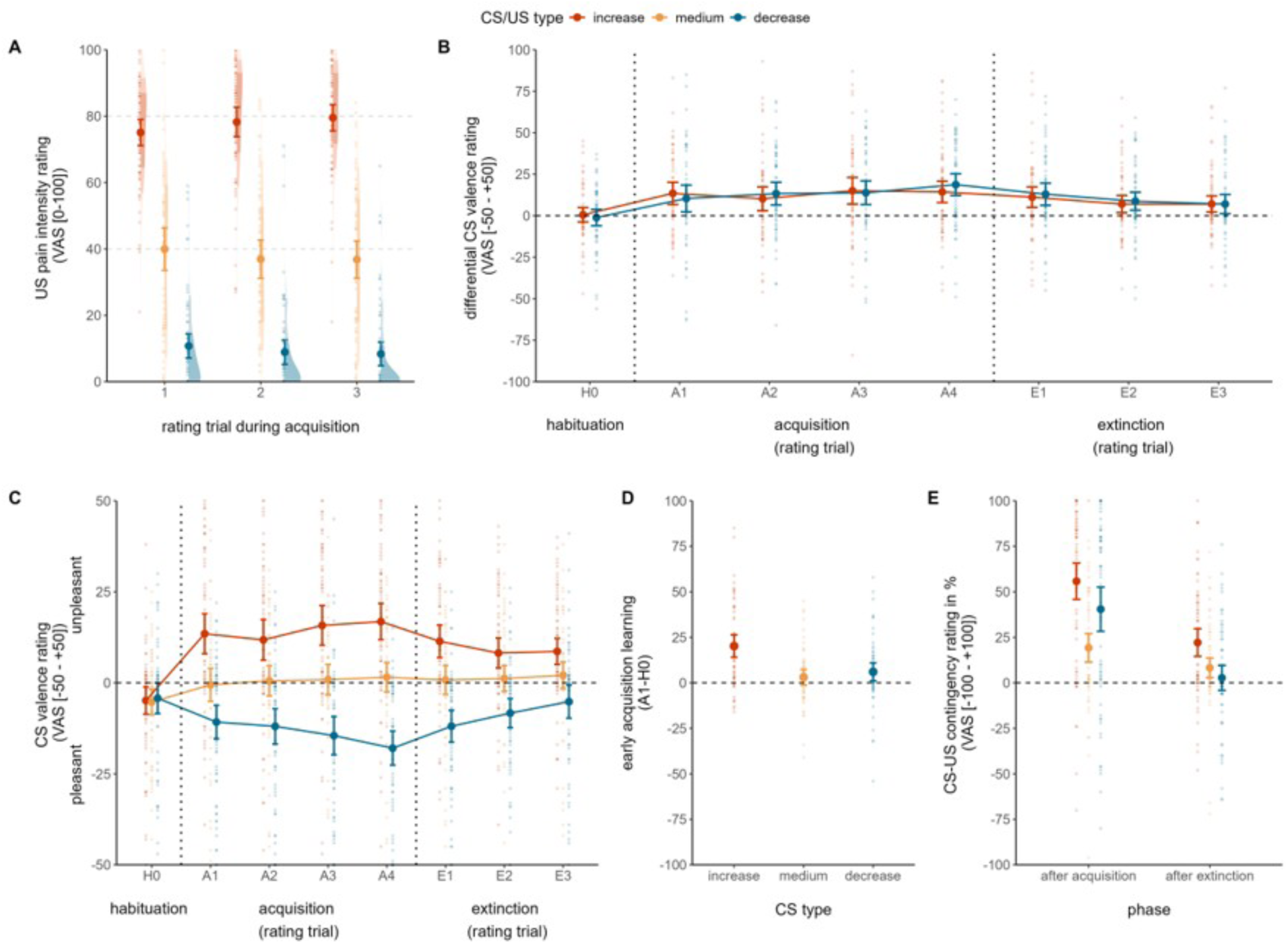
Behavioural results. Condition-wise means (dark-coloured dots) and 95% confidence intervals are displayed, along with individual data points (light colours). (A) US pain intensity ratings for the USincrease and the USdecrease were significantly different from the control condition (USmedium). (B) Differential valence ratings of CSincrease/decrease relative to the CSmedium did not differ between conditions during acquisition (A1-A4) or extinction (E1-E3). (C) Valence ratings of all three conditions (CSincrease, CSdecrease and CSmedium). Note that positive values indicate an unpleasant perception and negative values indicate a pleasant perception. (D) Early acquisition learning, defined as the difference in valence rating between the first acquisition rating (A1) and the habituation rating (H0), was most pronounced in the aversive (increase) condition. (E) CS-US contingency ratings, with positive values reflecting correct contingency learning for the CSincrease/decrease. CS = conditioned stimulus; US = unconditioned stimulus; VAS = visual analogue scale.

### Neuroimaging results

#### Acquisition phase: common neural responses during aversive and appetitive learning

To identify brain regions in which activation changed over time in both aversive and appetitive acquisition, we conducted a conjunction analysis ([CS_increase x time_ - CS_medium x time_] ν [CS_decrease x time_ - CS_medium x time_]). A joint activation increase was found in the occipital cortex, primarily encompassing visual areas V4 and V3 (Fig. 3A, Supplementary Table S4). The search for brain regions showing an activation decrease in both conditions yielded no significant result.

To further investigate these findings, we examined the functional connectivity of the two activated brain clusters during aversive and appetitive learning. Using the activated cluster in the occipital cortex as a seed region, we found increased positive coupling with the frontal operculum in both conditions (Fig. 3B; aversive: *t* = 4.21, *p*_SVC-FWE_ = .011, *k*_E_ = 7, MNI coordinates: [43,19,8]; appetitive: *t* = 3.80, *p*_SVC-FWE_ = .036, *k*_E_ = 5, MNI coordinates: [43,19,5]). This result suggests a synchronised increase in activity over time in the occipital cortex and frontal operculum, independent of CS valence.

#### Acquisition phase: distinct neural responses of aversive and appetitive learning

We also directly compared the appetitive and the aversive condition (relative to the CS_medium_ condition) to identify activations unique to each of them. As revealed by the interaction analysis ([CS_increase x time_ - CS_medium x time_] > [CS_decrease x time_ - CS_medium x time_]), the aversive condition induced a stronger signal increase in the right thalamus over the course of acquisition, specifically in the mediodorsal (MD) nucleus (Fig. 3C, Supplementary Table S4). No significant result was found for the opposite contrast testing for increasing activation in the appetitive condition ([CS_decrease x time_ - CS_medium x time_] > [CS_increase x time_ - CS_medium x time_]).

Using the identified cluster in the MD thalamus as a seed region, we investigated the functional connectivity of this cluster during differential learning over time ([CS_increase x time_ - CS_medium x time_] > [CS_decrease x time_ - CS_medium x time_]). This analysis revealed a stronger increase in positive connectivity with the right locus coeruleus (LC; Fig. 3D; *t* = 3.26, *p*_SVC-FWE_ = .018, *k*_E_ = 1, MNI coordinates: [6,-38,-23]), indicating a stronger increase in MD thalamus - LC coupling over the acquisition phase in the aversive than the appetitive condition.

#### Extinction phase: common neural responses of aversive and appetitive learning

To identify brain regions showing significant changes in activation over the course of extinction training in both conditions relative to the CS_medium_ condition, we conducted a conjunction analysis ([CS_increase x time_ - CS_medium x time_] ν [CS_decrease x time_ - CS_medium x time_]). While no cluster survived the threshold of *p*_SVC-FWE_ < .05, the vmPFC showed a trend-level decrease in response in both conditions (Fig. 4A, Supplementary Table S5). No regions exhibited shared increasing activities across the two conditions.

**Figure 3.**
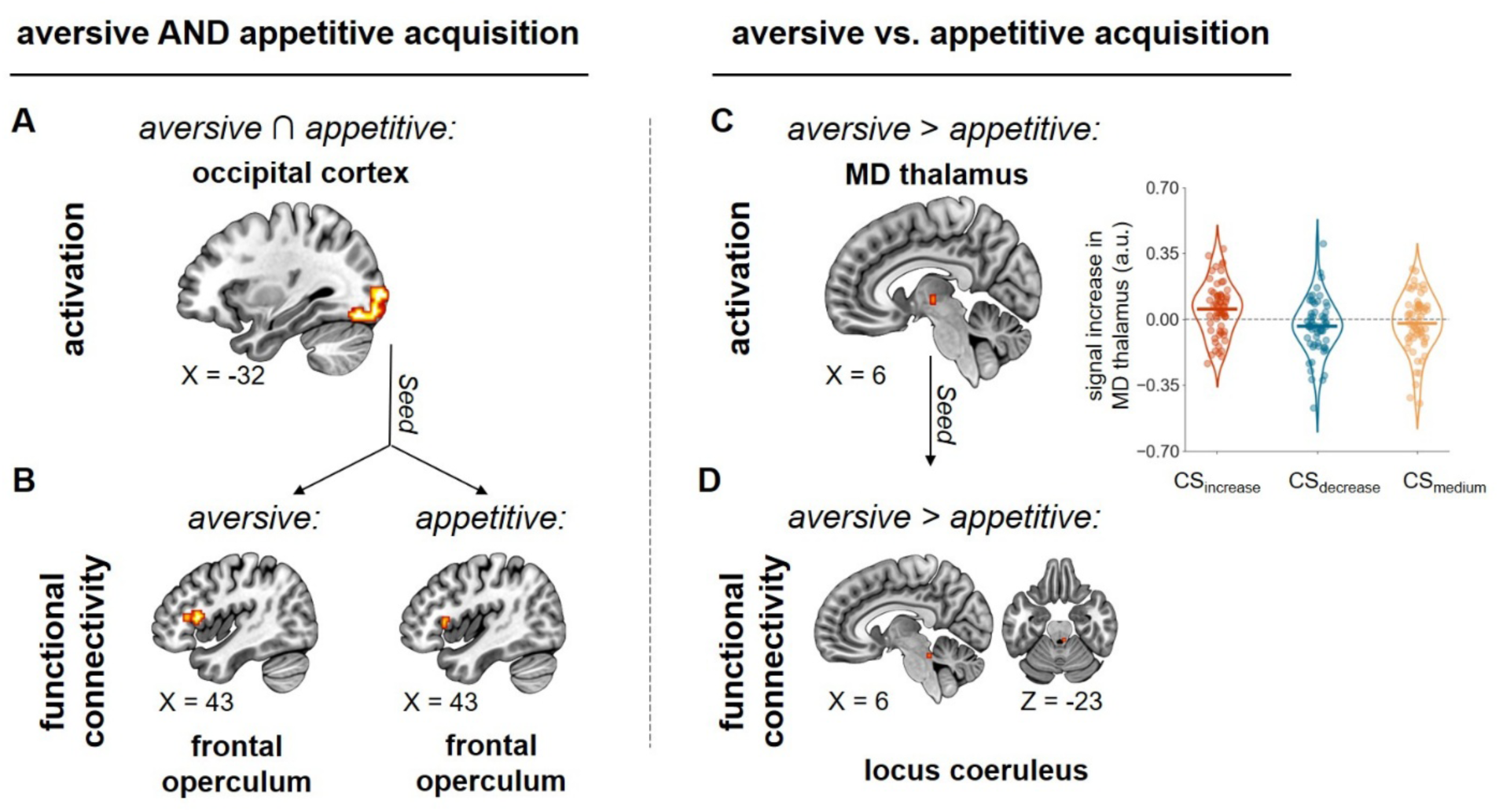
Joint and distinct activation and functional connectivity during CS events in the acquisition phase. (A) Brain regions showing a significant activation increase during both aversive and appetitive acquisition. (B) Functional connectivity of the cluster in the occipital cortex separately for both types of learning. (C) The right mediodorsal (MD) thalamus showed a stronger activation increase during aversive than appetitive acquisition. The inset displays parameter estimates (in arbitrary units) for each of the three conditions. (D) The cluster in the right MD showed a stronger increase in functional connectivity with the right LC during aversive than appetitive learning. a.u. = arbitrary units.

#### Extinction phase: distinct neural responses during aversive and appetitive learning

In the next step, we also compared the two conditions (both relative to the CS_medium_ condition). The contrast ([CS_decrease x time_ - CS_medium x time_] > [CS_increase x time_ - CS_medium x time_]) revealed a greater decrease in activation in the parahippocampal gyrus during extinction in the appetitive condition compared to the aversive condition (Fig. 4B, Supplementary Table S5). The reverse contrast ([CS_increase x time_ - CS_medium x time_] > [CS_decrease x time_ - CS_medium x time_]) yielded no significant result. To further investigate the differential engagement of the parahippocampal gyrus, we conducted a connectivity analysis using the identified cluster as a seed region. However, the search for brain regions showing a stronger decrease in functional connectivity with the parahippocampus during extinction showed no significant result.

**Figure 4.**
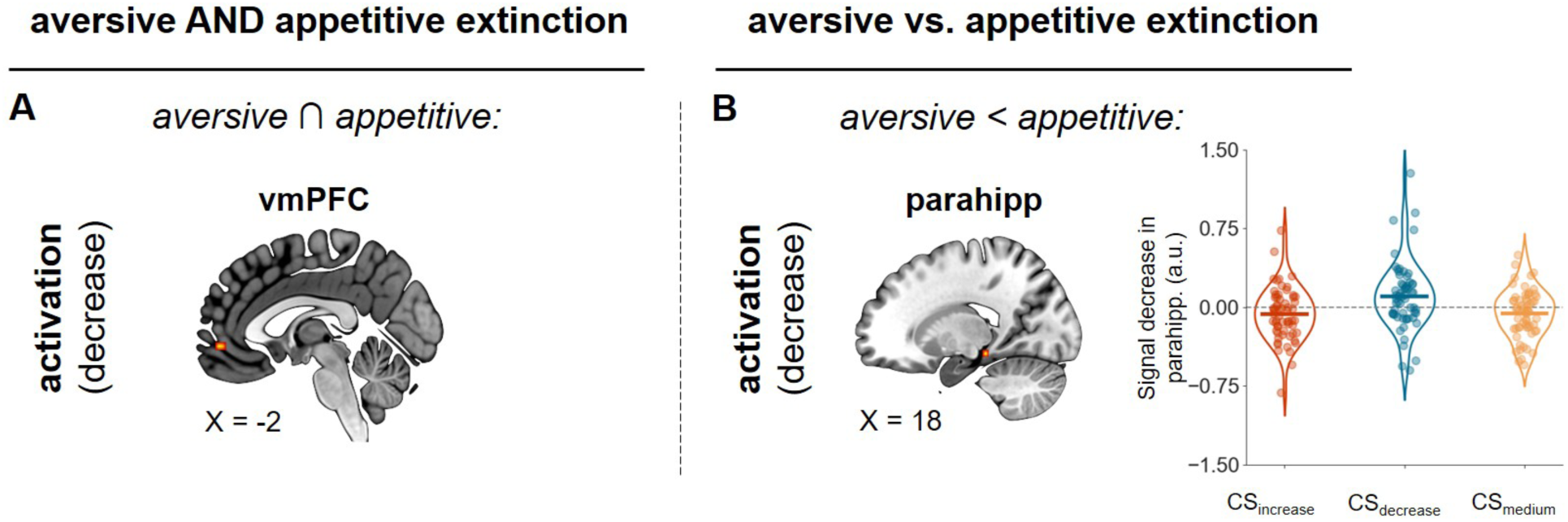
Joint and distinct activation and functional connectivity during CS events in the extinction phase. (A) The vmPFC showed a trend-level activation decrease during both aversive and appetitive extinction learning. (B) The parahippocampal gyrus displayed stronger activation decrease in the appetitive than the aversive condition. The inset displays parameter estimates (in arbitrary units) for each of the three conditions. vmPFC = ventromedial prefrontal cortex; parahipp = parahippocampal gyrus; a.u. = arbitrary units.

#### Residual anticipatory activity during the late CS phase as correlate of incomplete extinction

To investigate the neural basis of incomplete aversive extinction which we had also found previously^14^, we investigated activations during the late CS phase before US delivery as a proxy of (residual) pain anticipation^15^. To limit this search to brain regions involved in learning, we identified areas showing an activation increase during acquisition ([CS_increase x time_ – CS_medium x time_]) that also showed an activation decrease during extinction relative to the control condition ([CS_increase x time_ – CS_medium x time_]). During acquisition training, late CS presentation was accompanied by a signal increase in a left occipital cortex cluster at whole-brain correction level (Fig. 5A; *t* = 4.07, *p*_cluster-FWE_ = .029, *k*_E_ = 64, MNI coordinates: [-27,-93,-11]). Notably, the same occipital region also exhibited a signal decrease during the extinction phase (*t* = 4.25, *p*_SVC-FWE_ = .002, *k*_E_ = 4, MNI coordinates: [-32,-90,-17]), indicating its involvement in both learning phases. Parameter estimates extracted from this occipital cortex cluster showed a negative correlation with our behavioural index of incomplete extinction in the aversive condition (i.e., difference in contingency rating between CS_incease_ and CS_medium_ following extinction) (Fig. 5B; *r* = -.275, *p* = .032). This indicates that incomplete extinction in the aversive condition is linked to residual anticipatory engagement in the occipital lobe shortly before the US is presented. No significant result was found in other brain regions.

**Figure 5.**
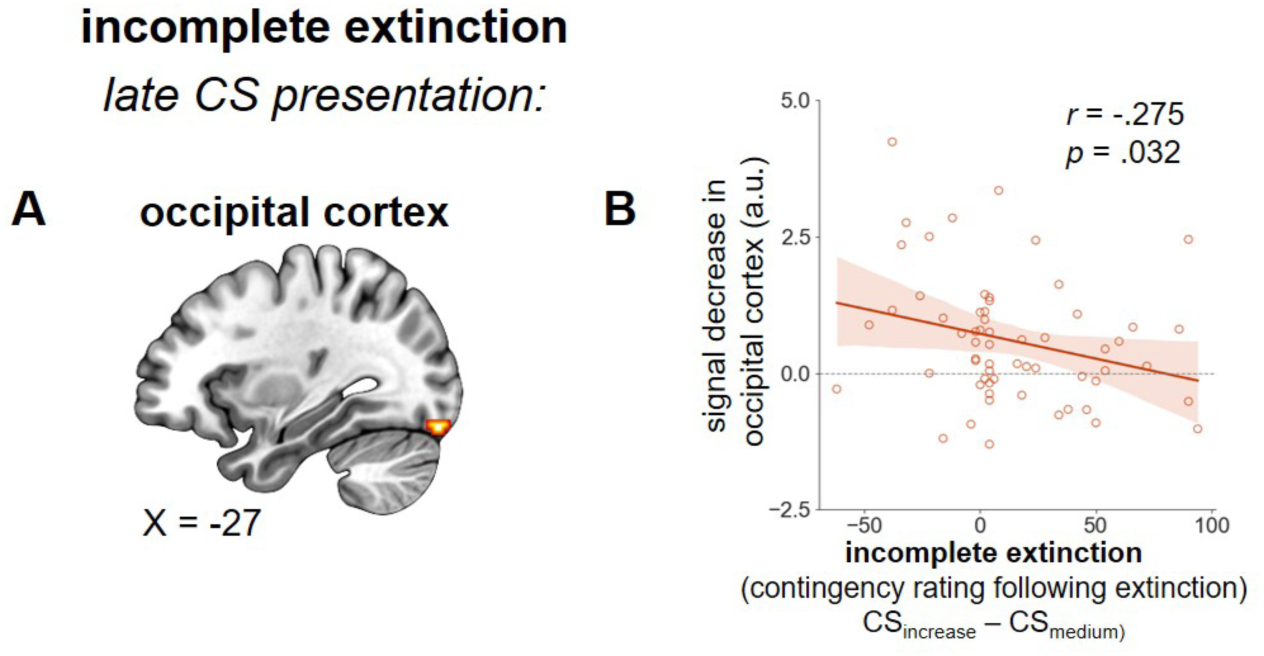
Occipital cluster involved in learning shows residual engagement during (anticipatory) late CS phase related to incomplete extinction. (A) The left occipital cortex showed an activation increase during acquisition [CSincrease x time - CSmedium x time] and an activation decrease during extinction in the late phase of aversive CS presentation [CSincrease x time - CSmedium x time], indicating its involvement in learning across phases. (B) The same occipital cluster showed residual activation in the late CS phase, with greater activation corresponding to more incomplete extinction in the aversive condition (as indicated by contingency ratings). CS = conditioned stimulus; a.u. = arbitrary units.

## Discussion

This study compared aversive and appetitive learning during acquisition and extinction on the behavioural and neural levels within the same paradigm. We report the following key findings: (1) A direct comparison of aversive and appetitive learning and early changes in valence ratings suggest stronger initial adjustments in aversive acquisition learning. Contingency ratings, reflecting cognitive aspects of learning, corroborate these findings indicating increased acquisition and incomplete extinction in the aversive condition. (2) Both types of learning led to an activation increase in the visual cortex during acquisition training and activity decrease in the vmPFC during extinction training. (3) During acquisition, aversive events induced a greater activity increase in the mediodorsal thalamus and functional connectivity with the locus coeruleus compared with appetitive events. (4) Appetitive extinction was accompanied by a greater activation decrease in the parahippocampal gyrus than extinction in the aversive condition. (5) Incomplete extinction in the aversive condition (as indexed by contingency ratings) scaled with residual activity in the visual cortex in the late CS phase linked to US anticipation.

As indicated by valence and contingency ratings, participants demonstrated enhanced acquisition learning for cues associated with pain exacerbation compared to pain relief. Moreover, initial learning was more pronounced for the CS_increase_, indicating fast adaptation to the aversive condition. But while valence ratings suggest that extinction was also comparable between both conditions, contingency ratings revealed a significant condition difference: At the end of the extinction phase, contingency ratings had returned to the pre-acquisition level in the appetitive condition, while they remained elevated in the aversive condition, suggesting incomplete extinction. This lingering memory for the more unpleasant outcome is in line with the so-called ‘better-safe-than-sorry’ strategy. Our findings thereby replicate a key result of the study by van der Schaaf & Schmidt et al. (2022) using the same paradigm^14^.

To better understand the neural mechanisms underlying these behavioural differences, we next examined joint and distinct brain activations during aversive and appetitive learning. Specifically, we investigated how neural activity and functional connectivity patterns evolve across acquisition and extinction, shedding light on the differential processing of aversive and appetitive cues.

During acquisition, both learning types showed significant signal increases in the occipital lobe, specifically within visual areas V3 and V4, in response to the conditioned stimulus. Notably, this occipital activation preceded the temperature modulation, in line with the concept of ‘preparatory attention’ - a mechanism that has been proposed to pre-activate neurons in the visual cortex relevant to enhance the perceptual sensitivity to anticipated stimuli^16^. As preparatory attention extends beyond spatial features to conceptual categories, it is considered content-based rather than feature-based^16^. Our finding of increased occipital activation during cue presentation in both aversive and appetitive acquisition supports this notion. In our paradigm, CS offset coincided with a change in visual input (Fig. 1B), shortly after the start in temperature change (i.e., increase or decrease). Since the display - a black screen - was identical across conditions, this change likely served as a general signal for an impending temperature shift rather than conveying specific directional information. The inferior frontal junction, including the operculum, has been implicated in driving preparatory attention in the visual cortex^17^, in line with our observations of increased functional connectivity between the occipital lobe and operculum during acquisition for both conditions (Fig. 3B). However, our data do not clarify the direction of information flow between these two regions.

When comparing aversive and appetitive acquisition learning, we observed that the right mediodorsal (MD) thalamus showed increased activation upon presentation of the visual cue signalling an increase in pain (Fig. 3C). The MD thalamus plays a crucial role in aversive learning through its connections with the amygdala and prefrontal regions, facilitating the integration of sensory and affective information essential for forming aversive memories that guide future behaviour and the execution and control of higher-level cognitive functions^18^. Additionally, the MD thalamus is well-connected to key attention-related regions, including the anterior cingulate cortex (ACC) supporting its role in prioritising relevant incoming information.

The stronger functional connectivity between the MD thalamus and the noradrenergic locus coeruleus (LC) over time during aversive learning provides additional evidence that the enhanced learning effect driven by aversive cues engages arousal and vigilance to potential threat and pain modulation^19,20^. The MD thalamus receives diverse inputs from brainstem structures, including the locus coeruleus (LC) and substantia nigra, with a topographical organisation^21^. LC is a main source of noradrenaline that modulates alertness and arousal. The increased connectivity between both regions might therefore not only reflect amplified attention to the aversive increase in pain but also prepare for adaptive responses, such as avoidance and withdrawal behaviour.

During extinction, both aversive and appetitive extinction shared a trend-level signal decrease over time in the vmPFC (Fig. 4A), a key region widely implicated in fear extinction across humans and animal studies^22–24^. Notably, extinction is temporally sensitive^25^ and as Morriss and colleagues^26^ commented, a more statistically powerful approach should be employed such as applying time modulators to regressors to capture linear changes over time^27,28^. Following these suggestions, we involved time-based comparisons and observed temporal dynamics in the vmPFC independent of the valence of extinction.

The direct comparison of appetitive and aversive extinction learning revealed a weaker signal decrease in the right parahippocampal gyrus (PHG) during the aversive condition (Fig. 4B). This pattern may reflect the PHG’s role in encoding and retrieving contextual associations, with notable differences in how appetitive and aversive memories are processed. For appetitive stimuli, which are often less critical for survival, the brain may more easily disengage from positive associations, as reflected in a stronger PHG deactivation during extinction. In contrast, aversive stimuli often sustain a ‘safety monitoring’ component due to their evolutionary significance, maintaining a heightened vigilance in the PHG even as extinction progresses. Together, these findings suggest that the PHG’s decreased activation during appetitive extinction reflects the brain’s readiness to release positive associations, while its sustained activation and connectivity with regulatory and sensory regions during aversive extinction highlight a cautious approach, preserving threat awareness and adaptive responses. This may explain the brain’s tendency to ‘let go’ of rewarding cues more readily than those linked to potential threats.

To further explore the neural underpinnings of incomplete extinction suggested by contingency ratings in more detail, we conducted an additional analysis focusing specifically on the late phase of the conditioned stimulus (CS) presentation, a period when participants begin anticipating the unconditioned stimulus (US), as suggested by Eippert and colleagues^15^. This approach allows us to capture anticipatory processes that may differ between appetitive and aversive conditions. If extinction learning in the aversive condition is indeed incomplete—as indicated by our contingency data analysis—we would expect sustained activation in regions associated with pain anticipation, reflecting the persistence of conditioned threat responses. The analysis revealed that incomplete extinction was related to a lesser decrease in occipital cortex activity during extinction (Fig. 5). In line with the notion of preparatory attention discussed above, the weaker signal decrease in the visual cortex during extinction in the aversive relative to the neutral condition may reflect a sustained state of preparatory attention in response to the aversive stimulus.

In summary, we demonstrate that aversive and appetitive learning share mechanisms but also rely on distinct processes involving pain-related, learning-associated, and preparatory attention regions. Aversive learning uniquely engages areas promoting rapid acquisition such as the mediodorsal thalamus and locus coeruleus. Importantly, we provide converging evidence for incomplete extinction, primarily in the occipital cortex but also in the parahippocampal gyrus. The weaker signal decrease in the PHG during aversive extinction suggests a sustained engagement of contextual memory processes, likely reflecting an ongoing monitoring of potential threats. Similarly, residual activity in the occipital cortex during the late CS phase aligns with a persistent state of preparatory attention, maintaining heightened sensitivity to aversive cues. Together, these findings highlight distinct neural mechanisms underlying aversive extinction, where threat-related associations are more resistant to suppression, contributing to the observed behavioural evidence of incomplete extinction. Such ‘better-safe-than-sorry’ strategy is discussed as a key factor in the development and maintenance of pain and other conditions related to aberrant learning such as anxiety disorders.

## Acknowledgements

We would like to thank Matthias Gamer and Balint Kincses for conceptual and methodological support. This study was funded by the German Research Foundation—project A11, project number 316803389—SFB 1280 (Gefördert durch die Deutsche Forschungsgemeinschaft (DFG)— Projekt A11, Projektnummer 316803389—SFB 1280).

## Methods

### Participants

A total of *N* = 84 healthy, right-handed participants were recruited through advertisements. Exclusion criteria were age < 18 or > 75 years, a self-reported history of acute or chronic pain or other neurological or psychiatric disorders, regular use of medication (including hormonal contraception), contraindication to MRI scans, pregnancy or breastfeeding, allergy to capsaicin, acute sunburn or other dermatological abnormalities on the volar forearm, BMI > 30 or < 18, and left-handedness. The study was registered at the German register for clinical trials (https://www.drks.de/search/de/trial/DRKS00027448/details) and approved by the local Ethics Committee (16-7248-BO) of the Medical Faculty, University Duisburg-Essen. Participants gave written informed consent before the start of the experiment and received a monetary compensation for their participation (50 Euro plus 15 Euro/hour). Participants could withdraw from the study at any time.

### Experimental procedures and paradigm

The study was conducted over two experimental days (mean number of interval days = 2.31, range [0, 4]). On day 1, participants underwent a calibration procedure to determine the individual temperature levels for the thermal stimuli. Moderate continuous pain was elicited using a combined capsaicin and tonic heat model^12,14^. Capsaicin cream (1%, 8-methyl-Nvanillyl-6-nonenamide, 98%, Sigma, diluted in 5% ethanol-KY jelly) was applied to the volar forearm over an area of 3x3cm. Capsaicin is an active ingredient of chili peppers that increases thermal sensitivity by binding to vanilloid receptors, which allows low-level heat stimuli to evoke tonic heat pain that mimics clinical pain with lasting effects through controllable temperature manipulations^29^. After approximately 45 min, during which participants completed a battery of questionnaires to collect demographic information and assess psychological variables (details see Questionnaires) the capsaicin cream was removed and thermal stimulation (Model ATS, Pathway System, Medoc, Israel) was applied to the capsaicin pre-treated area. The individual heat pain threshold was then determined using a previously validated temperature calibration procedure^14^ (for details see Supplementary Methods).

On day 2, participants completed additional questionnaires (details see Questionnaires) after the capsaicin cream was again applied. Approximately 45 min later, participants were positioned in the MRI scanner and their heat pain thresholds were reassessed (for details see Supplementary Methods). Participants then completed the differential conditioning task in the scanner (for details see Differential conditioning paradigm), which was followed by high-resolution whole-brain T1-weighted and DTI scans. Note that we only report the task-based fMRI results here, and results from the DTI scans will be reported elsewhere.

### Differential conditioning paradigm

Participants underwent a differential conditioning paradigm comprising a habituation, acquisition and extinction learning phase. During both the acquisition and the extinction phases, individually calibrated contact heat stimuli (unconditioned stimuli, US) were paired with predictive cues (conditioned stimuli, CS) (Fig. 1). Moderate heat stimuli were continuously applied to induce the ongoing tonic pain throughout the experiment. The CS-US association was examined in three experimental phases: habituation (CS familiarization), acquisition training (CS presentation with a 75% US reinforcement rate) and extinction training (CS presentation without reinforcement). Three blue geometric figures (a square, rectangle and rhombus; RBG: 142,180,227; visual angles: square 4.99° × 4.99°, rectangle 8.3° × 3.14°, rhombus 7.38° × 5.36°) served as CS (CS_increase_, CS_decrease_, CS_medium_) to predict pain exacerbation (US_increase_), pain decrease (US_decrease_) and no change in temperature (US_medium_), respectively. The assignment of the geometric figures to the CS conditions was balanced and pseudo-randomized across participants.

*Habituation*. Prior to the experiment, participants were instructed about the presentation of the geometric cues during habituation and were informed about the potential CS-US association in the following phases, though they were not given details about the actual contingency. During habituation, each CS was presented three times without any change in temperature (US_medium_).

*Acquisition training*. During the acquisition training, a total of 48 CS was presented (16 per type). The CS_increase_ and CS_decrease_ were each paired with their corresponding US (US_increase_ and US_decrease_) at a 75% reinforcement rate, resulting in 12 reinforced CS per type. The CS_medium_ was followed by no change in temperature (US_medium_). The first and last trials of each condition were always reinforced.

*Extinction training*. Following the acquisition training, no additional instructions were provided. During extinction training, a total of 36 CS (12 per type) were presented with US_medium_ only (i.e., ongoing thermal stimulation with temperature corresponding to VAS40).

*Trial structure.* On each trial, the CS was presented for 9s, followed by a black screen for 10.5s. The US were applied for 8s, starting 1.5s before the offset of the CS (i.e., CS and US overlapped for 1.5s), with a 6.5s overlap with the presentation of the black screen. A white fixation cross was displayed during the inter-trial-intervals (ITI) that were jittered between 6-11s. During both acquisition and extinction training, CS were displayed in a pseudo-randomized order, ensuring that (1) the same CS type was presented no more than three times consecutively and (2) each CS type was equally distributed across both halves of the experimental phases.

The software Presentation (www.neurobs.com) was used to present visual stimuli, trigger application of thermal stimuli and record behavioural data.

### Behavioural outcome measures

To track the temporal dynamics of the acquisition and extinction of CS-US associations at the emotional as well as the cognitive level^13,14^, participants provided ratings for CS valence (“How do you perceive this geometric figure?”) and CS-US contingency (“How often was this geometric figure followed by a painful stimulus?”) using a Visual Analogue Scale (VAS). The valence scale was anchored at -50 = “very pleasant”, 0 = “neutral” and 50 = “very unpleasant”. The conÉngency scale was anchored at -100 = “100% cooling”, 0 = “no change” and 100 = “100% heaÉng”. Note that the numerical values of the anchors were never visible to the participants. The VAS was displayed for 7s, beginning at the onset of the corresponding CS presentation. During the habituation phase, participants provided one valence rating for each CS type at the beginning of the phase. Further valence ratings were collected on every fourth CS presentation during acquisition training (four ratings per CS type) and extinction training (three ratings per CS type). CS-US contingency ratings were given at the end of both acquisition and extinction training, with no time limit imposed.

Ratings of US pain intensity and (un)pleasantness during acquisition and extinction training were collected using VASs (US pain intensity: “How painful was this temperature stimulus”; anchors: 0 = “not painful at all” and 100 = “unbearably painful”; US (un)pleasantness: “How pleasant/unpleasant was this temperature stimulus?”; anchors: −50 = “very pleasant”, 0 = “neutral” and 50 = “very unpleasant”). Following every fourth US of the same type, participants were given 15s to rate each question, with an ITI of 0.5s between questions. After confirming their responses for US pain intensity, the fixation cross of the ITI was immediately displayed, followed by the US (un)pleasantness rating (i.e., duration of US ratings: individual reaction time). This resulted in three ratings for each US type during acquisition training and five ratings for US_medium_ during extinction training. Prior to habituation, participants rated their current arousal level (“How tense do you feel at the moment?”, anchors: from “not tense at all” to “extremely tense”) and pain-related fear (“How fearful are you about the upcoming pain sÉmulaÉon?”, anchors: from “not fearful at all” to “extremely fearful”) using VAS 0-100. For each rating, the VAS cursor was placed at a random starting position between VAS25 and VAS75. Participants were instructed to confirm their responses and the confirmed ratings were used for behavioural analyses. Except for contingency and US ratings, the VAS presentation remained on screen after confirmation until the next event started. A summary of the number of presentations and ratings can be found in Supplementary Table S3.

### Questionnaires

Participants completed the German version of the following psychological questionnaires: State Trait Anxiety Depression Inventory: STADI^30^; Pain Anxiety Symptom Scale: PASS-20^31^; Pain Catastrophizing Scale: PCS^32^; Perceived Stress QuesÉonnaire: PSQ-20^33^; and Depression Anxiety Stress Scales: DASS^34^. Participants completed the STADI-state on day 2, and STADI-trait and the remaining questionnaires on day 1. All questionnaires were analysed according to their respective manuals. The questionnaire results can be seen in Supplementary Table S1.

### Skin conductance

Due to technical issues, SCR data from *n* = 41 participants had to be excluded, leaving a dataset of only *n* = 27. Given the substantial data loss, we decided not to report these analyses.

### Behavioural analyses

Behavioural analyses of CS valence ratings, contingency ratings, US intensity and (un)pleasantness ratings were carried out in R version 4.1.1^35^. To compare aversive and appetitive acquisition and extinction learning linear mixed models (LMM) were estimated using the lme4 package^36^ (version 1.1-31) for each learning phase separately. In general, models were estimated using the restricted maximum likelihood (REML) approach and always included the factors specified in the model design as fixed effects (see below for the individual models and Supplementary Table S2). First, significant model improvement was tested by including a random intercept by participant. Second, improvement was tested by adding random slopes by CS (or US) type. All model comparisons were assessed using likelihood ratio tests and a reduction in the Akaike’s information criterion (AIC). If the addition of a factor significantly improved the model fit, the more complex LMM was used. The significance of the included factors, as well as their interactions, were assessed using Type III Wald chi-square tests. We report beta values from the model results when they are sufficient for interpretation (i.e., in the case of significant main effects involving only the respective factors). For specific differences in significant interaction effects, we report post-hoc tests carried out using the emmeans package^37^ (version 1.8.3). A correction for multiple testing was applied using the Tukey or Sidak method where appropriate.

#### US intensity and US (un)pleasantness

Ratings of US pain intensity and US (un)pleasantness were analysed using LMMs to ensure that the calibration procedure was successful and that the three stimulation temperatures implemented as US_increase_, US_medium_ and US_decrease_ yielded different pain intensities and (un)pleasantness levels during the acquisition phase. For details on analyses and results see Supplementary Information.

#### CS valence

To assess acquisition and extinction learning on the behavioural level, CS valence ratings were analysed using a LMM with participant as a random effect (random intercepts, and random slopes by *CS type*), and *CS type* and *time* (continuous), as well as their interaction, as fixed effects. CS valence ratings were first analysed differentially, i.e., relative to the CS_medium_ (i.e., CS_increase_ - CS_medium_; CS_medium_ - CS_decrease_), to directly compare the level of CS-US associations between the increase and decrease conditions. In a second model, ratings for each of the three conditions were entered. Models were estimated separately for each phase, using the final rating from the previous phase as the baseline (i.e., H0 – A4 for acquisition training and A4 – E3 for extinction training; see also Fig. 2). Models included fixed effects of *CS type* (CS_increase_, CS_medium_, and CS_decrease_, or CS_increase_ and CS_decrease_ (relative to the CS_medium_)) as a categorical factor and *time* as a continuous variable, as well as their interaction. Additionally, we conducted an exploratory comparison of the aversive and appetitive conditions using paired t-tests on individual slopes.

#### CS-US contingency

To investigate acquisition and extinction learning on the cognitive level, we analysed CS-US contingency ratings provided at the end of each learning phase. CS-US contingency ratings were transformed so that positive values represented correct learning (i.e., CS_decrease_ value x (-1)). Models included the same fixed effects factors as described for CS valence, including all three levels of *CS types* (i.e., CS_increase_, CS_medium_, CS_decrease_), except that *time* was added as a categorical factor (i.e., rating after acquisition, and after extinction training only). To test for incomplete extinction for the aversive or appetitive condition, CS-US contingency ratings were compared against 0 by phase and CS type using post-hoc comparisons.

### fMRI data acquisition and analysis

MRI data were collected on a 3.0 Tesla scanner (Siemens Trio) with a 32-channel head coil. Functional images were obtained using a gradient EPI T2*-weighted sequence for each participant with the following acquisition parameters: repetition time (TR), 2.19s; echo time (TE), 26ms; slices number, 42; slice-thickness, 2.5mm; field of view (FOV), 224×224mm; voxel size, 2.5×2.5×2.5mm; flip angle, 90°. High-resolution whole-brain T1-weighted images were acquired using an MP-RAGE sequence with TR, 1.77s; TE, 3.24ms; slices number, 224; slice-thickness, 1mm; FOV, 256×256mm; flip angle, 8°.

#### Data preprocessing

Neuroimaging data were preprocessed using the fMRIprep v22.1.1 pipeline^38^. Preprocessing of participants’ T1-weighted images included skull-stripping, spatial normalization and brain tissue segmentation. Functional images were preprocessed separately for acquisition and extinction training, including co-registration with individual T1-weighted reference image, motion correction, slice-timing correction and normalization to MNI152NLin2009cAsym standard space. In each phase, physiological regressors were calculated using component-based noise correction (CompCor)^39^, and the first 6 anatomical components (aCompCor) for each participant were added as nuisance regressors to the individual design matrix in the first-level analysis. Functional images were spatially smoothed using a Gaussian kernel with 6mm full-width at half-maximum (FWHM).

#### fMRI data analysis

Univariate general linear model (GLM) analyses were conducted separately for the acquisition and extinction phases using SPM12 (https://www.fil.ion.ucl.ac.uk/spm/). To perform all planned analyses, the individual design matrices (subject level) included six regressors of interest coding for the CS events (CS_increase_, CS_decrease_, CS_medium_). CS phases were segmented into an early response, starting at CS onset, and a late response, starting 4.5s after CS onset (modelled duration for each event: 0s) adopting the approach described by Eippert and colleagues^15^ to investigate potential incomplete extinction, as found in our previous study^14^. The late response during the CS phase is thought to reflect US anticipation, which, in the case of incomplete extinction, would be expected to remain elevated (i.e., show less decrease over time). To examine brain activity during pain exacerbation and pain relief in the acquisition phase, two regressors modelling US_increase_ and US_decrease_ (duration: 8s) were added to the individual design matrices). For both phases, the individual design matrices also included nuisance variables controlling for motion-related artefacts, including head movement (6 aCompCor components) and ratings (valence: duration 7s; US pain intensity and (un)pleasantness: individual rating time as duration with a maximum of 15s per question). Both CS and US events were convolved with a canonical hemodynamic response function. To account for learning over the course of each phase, we applied time modulation (linear increase) to the CS regressors in each model as done in previous studies^15,28^.

Individual-level contrasts for time-modulated CS events (combining early and late CS responses: [CS_increase x time_ - CS_medium x time_], [CS_decrease x time_ - CS_medium x time_]) and the beta images modelling individual US conditions (acquisition training: US_increase_, US_decrease_) were included in group-level analyses. Individual-level contrasts for CS events without time modulation (i.e., average activity over the course of each learning phase) were used for supplementary analyses.

The following analyses were performed to answer our research questions: (1) Common mechanisms underlying aversive and appetitive learning were examined using a conjunction analysis ([CS_increase x time_ - CS_medium x time_] ∩ [CS_decrease x time_ - CS_medium x time_]), conducted separately for the acquisition and extinction phase^40^. (2) To identify differences between aversive and appetitive acquisition and extinction learning as well as between phases of pain exacerbation and pain relief, differential contrasts ([CS_increase x time_ - CS_medium x time_] > [CS_decrease x time_ - CS_medium x time_], [US_increase_ > US_decrease_]) were calculated separately for CS and US events and entered into group-level one-sample *t* tests. Note that, in line with aims of our primary analysis, the main results in the study are based on activity changes over time during CS events. In addition to these time modulation effects, the overall average activity during the acquisition and extinction phases was calculated using the contrasts images without time modulation (e.g., [CS_increase_ - CS_medium_] > [CS_decrease_ - CS_medium_], [US_increase_ > US_decrease_]). Furthermore, neural responses to aversive and appetitive learning over the course of acquisition and extinction training were analysed (e.g. CS_increase x time_ > CS_medium x time_). Results of these supplementary analyses are provided in the Supplementary Information. (3) To further explore the role of the brain regions identified in the activation analyses (e.g., the areas sharing activity increase over time during both types of learning), we investigated the functional connectivity of these regions using gPPI^41^ separately for the acquisition phase and the extinction phase. The corresponding first-level models comprised psychological variables (the same regressors as in the BOLD activation design matrix), physiological variables (time series from seed region) and the respective psychophysiological interaction terms (multiplication of psychological and physiological variables). The time modulation effect was applied to the interaction terms in the design matrix, allowing us to test how the strength of functional connectivity between the seed region and other regions changes over time in relation to each condition. A significant positive connectivity result indicates that the increasing activities in the seed region and the connected region are positively correlated over time. For the seed regions which showed shared activity changes over time during both conditions, we examined functional connectivity separately for appetitive and aversive learning using the corresponding contrasts ([CS_decrease x time_ - CS_medium x time_], [CS_increase x time_ - CS_medium x time_]). We also performed seed-to-voxel analyses on differential learning using the differential contrasts ([CS_increase x time_ - CS_medium x time_] > [CS_decrease x time_ - CS_medium x time_]). Individual-level contrast images were entered into group-level one-sample *t*-tests. (4) To explore neural processes underlying the incomplete extinction we found on the behavioural level (i.e., elevated contingency ratings for the CS_increase_ after extinction training; see Results), we took a 2-step approach. To focus on brain regions reflecting learning, we first identified brain regions exhibiting signal increase in the late CS phase during acquisition training using the contrast [CS_increase x time_ – CS_medium x time_], with the occipital cortex emerging as the only cluster surviving whole-brain correction. Using this occipital cluster as a region of interest (ROI), we then observed a signal decrease in the late CS phase during extinction training (*p*_SVC-FWE_ < .05). In the second step, activation change during extinction training in this cluster was correlated with our behavioural measure of incomplete extinction (contingency rating of CS_increase_ – CS_medium_ after extinction). The BOLD signal from the activated cluster was extracted using MarsBaR^42^.

We used a region-of-interest approach on brain networks involved in pain, learning, reward processing and salience^3,8,43–45^. ROIs included the OFC, vmPFC, dmPFC, frontal operculum, insula, S1, S2, parahippocampal gyrus, hippocampus, amygdala, striatum, VTA, and brainstem. The brainstem ROI (i.e., locus coeruleus) was defined using probabilistic anatomical grey matter masks^46^. All other ROIs were defined using anatomical masks based on Harvard-Oxford cortical and subcortical 25% probability atlases^47^. Within these regions, results were thresholded at *p* < .05 family-wise error (FWE) corrected at peak level with small volume correction (SVC). Outside these regions, we report results at the whole-brain level (*p*_cluster-FWE_ < .05). Subregion identification for the occipital cortex was based on Julich-Brain Cytoarchitectonic Atlas^48^, and The Thalamus Atlas^49^ for the thalamus.

## Notes

### Competing Interest Statement

The authors have declared no competing interest.

