## Supplementary Information for "Common and distinct neural mechanisms of aversive and appetitive pain-related learning"

### Supplementary methods

#### Temperature calibration

To experimentally induce tonic heat pain and take into account the sensitization and habituation of pain perception that have been reported in previous studies[1,2], a temperature multi-step calibration procedure was performed for each participant on both experimental days. Individual pain perception was assessed using Visual Analog Scale ratings of pain intensity (VAS 0-100 with anchors 0 = "not painful at all" and 100 = "unbearably painful"). The below described process yielded three temperature levels that were perceived as intensely painful (VAS 80), moderately painful (VAS 40), and not painful (VAS 0), which were used for $US_{increase}$, $US_{medium}$ and $US_{decrease}$, respectively, throughout the experiment.

On day 1, we first applied a ***staircase*** procedure twice with the temperature manipulation starting at 28°C and an increase of 2°C per step (when reaching 42°C: 1°C rise for each step; maximum temperature: 47°C) until participants reported a VAS = ~60. The temperature of VAS 60 was then used to determine the ten temperature levels (ranging from -1.5°C to +3°C, with a step of 0.5°C) in the ***regression*** session, where each level was applied twice in a semi-randomized order. The temperature remained constant for 8s before returning to baseline level (26°C). Temperature levels of VAS 80 and VAS 60 were estimated by linear regression in R. The temperature level for the $US_{decrease}$ was calculated by the level corresponding to VAS 60 minus approximately 10°C (minimum temperature level applied: 20°C) to achieve a non-painful, relieving sensation of VAS 0. In order to account for habituation effects, which particularly affect the tonic moderately painful stimulus[3], the temperature corresponding to a VAS60 as determined in the regression session was used to induce a lasting sensation corresponding to a VAS40 ($US_{medium}$). This was then tested by applying this temperature continuously and obtaining four pain intensity and unpleasantness ratings from the participants. The subsequent ***training*** session started with the continuous application of the VAS 40 temperature, followed by three presentations for each of the temperature levels for $US_{increase}$ and $US_{decrease}$ in one of six predefined orders. For each temperature level, participants were instructed to rate pain intensity and (un)pleasantness using VAS (for US pain intensity: 0 = "not painful at all" and 100 = "unbearably painful"; for US (un)pleasantness: −50 = "very pleasant", 0 = "neutral" and 50 = "very unpleasant") for three times. Based on these pain intensity ratings the stable temperature levels corresponding to VAS 80, VAS 40 and VAS 0 were determined and used for $US_{increase}$, $US_{medium}$ and $US_{decrease}$, respectively.

On day 2, we reassessed the three temperature levels by repeating the *regression* and *training* sessions in the scanner. The calibrated three temperature levels were employed for the corresponding US in the experimental phases.

**Behavioural covariate analyses**

The following psychological variables were entered as covariates in separate analyses: arousal and pain-related fear ratings, and two composites of aggregated questionnaire scores, one for affective measures (STADI trait anxiety and depression scales and the global score of the PSQ-20 [stress]), and one for pain-related measures (PCS [pain catastrophizing] and PASS-20 [pain anxiety] global scores). Raw scores of each scale were *z*-standardised across participants, and the resulting scores were averaged for each composite score and each participant. The respective factors were separately added to the different models (i.e., CS valence during acquisition, CS valence during extinction etc.) which were compared to the basic model using likelihood ratio tests and a reduction in the Akaike's information criterion (AIC). If the addition of a factor significantly improved the model fit, we report differences compared to the basic model.

**fMRI data analyses**

***Time-modulated neural activity***

Beyond our primary results, to facilitate future meta-analysis studies, we additionally report neural responses to aversive and appetitive learning over the course of acquisition (see *fMRI result 1* and Supplementary Table S6) and extinction training (see *fMRI result 2* and Supplementary Table S7) separately. Individual contrasts with $CS_{medium\ x\ time}$ ([$CS_{increase\ x\ time}$ > $CS_{medium\ x\ time}$], [$CS_{decrease\ x\ time}$ > $CS_{medium\ x\ time}$]) were calculated and entered into group-level one-sample *t* tests. Results were thresholded at $p_{uncorrected} < .001$ at peak-level.

***Average neural activity***

In addition to time modulation effects, the overall average activity within each training phase was calculated (i.e., GLM without time modulators). Common and distinct mechanisms underlying aversive and appetitive conditions were investigated separately for both CS events during the acquisition (see *fMRI result 3* and Supplementary Table S8) and extinction training (see *fMRI result 4* and Supplementary Table S9), and US events (see *fMRI result 5* and Supplementary Table S10). Individual contrasts (e.g., [$CS_{increase}$ - $CS_{medium}$] $\cap$ [$CS_{decrease}$ - $CS_{medium}$], ([$CS_{increase}$ - $CS_{medium}$] > [$CS_{decrease}$ - $CS_{medium}$]), [$US_{increase}$ > $US_{decrease}$]) were calculated and entered into group-level one-sample *t* tests. Results were thresholded at $p_{uncorrected} < .001$ at peak-level.

### Supplementary results

#### US pain intensity and US (un-)pleasantness

##### *Difference in US pain intensity perception between US types during acquisition training*

During the acquisition training, US pain intensity ratings differed significantly between *US types* ($X^2(2) = 235.17$, $p < .001$, post-hoc tests: $US_{increase} > US_{medium}$ $\Delta\beta = 39.70$, $t(67) = 17.24$, $p < .001$; $US_{medium} > US_{decrease}$ $\Delta\beta = 28.50$, $t(67) = 12.52$, $p < .001$). This indicates that overall, the calibration procedure was successful. The main effect of *time* was not significant ($X^2(1) = 1.73$, $p = .188$), while the *US type* x *time* interaction was significant ($X^2(2) = 6.07$, $p = .048$). However, slopes did neither differ between *US types* ($US_{increase}$ - $US_{medium}$ $\Delta\beta = 3.79$, $t(404) = 2.26$, $p = .064$; $US_{medium}$ - $US_{decrease}$ $\Delta\beta = -0.45$, $t(405) = -0.27$, $p = .962$, $US_{increase}$ - $US_{decrease}$ $\Delta\beta = 3.34$, $t(405) = 1.98$, $p = .118$), nor did they differ from zero ($CS_{increase}$ $\beta = 2.23$, $t(404) = 1.87$, $p = .174$; $CS_{medium}$ $\beta = -1.57$, $t(404) = -1.32$, $p = .466$; $CS_{decrease}$ $\beta = -1.12$, $t(406) = -0.93$, $p = .726$). Thus, there was no significant change in individual US perception over time, as calibrated differences lasted during the acquisition training.

##### *Difference in US (un)pleasantness perception between US types during acquisition training*

A similar pattern as for US pain intensity emerged for US (un)pleasantness (see Fig. S1), which differed by *US type* ($X^2(2) = 142.97$, $p < .001$, post-hoc tests: $US_{increase} > US_{medium}$, $\Delta\beta = 25.80$, $t(67) = 14.17$, $p < .001$; $US_{medium} > US_{decrease}$, $\Delta\beta = 34.70$, $t(67) = 14.64$, $p < .001$) with higher ratings for the $US_{increase}$ than the $US_{medium}$, and lowest (i.e., most pleasant) ratings for the $US_{decrease}$. US (un)pleasantness ratings did not change significantly over time (main effect *time* $X^2(1) = 0.91$, $p = .340$; interaction *US type* x *time* $X^2(2) = 5.06$, $p = .080$).

##### *Changes in US pain intensity and US (un)pleasantness for the US medium*

Changes in mean pain intensity and mean pain (un)pleasantness of the tonic pain stimulus ($US_{medium}$) from acquisition to extinction training were investigated using separate paired t-tests. Both analyses indicated that participants habituated to the tonic painful stimulation during the experimental phases. More specifically, pain intensity was significantly lower during the extinction phase than during the acquisition phase ($t(67) = 5.17$, $p < .001$; acquisition: M = 37.88, SD = 19.98; extinction: $M = 27.41$, $SD = 17.40$). Similarly, pain (un)pleasantness rating were significantly lower (i.e., more pleasant; $t(67) = 3.96$, $p < .001$) in the extinction phase ($M = -1.88$, $SD = 17.22$) than in the acquisition phase ($M = 5.15$, $SD = 13.91$).

**Influence of covariates of interest on behavioural analyses**

We also tested whether including arousal, pain-related fear, or the two composite scores (general affective and pain-related) to all of the models reported in the main analyses (i.e., acquisition and extinction phase; CS valence and CS-US contingency ratings; differential and non-differential) improved model fit; however, none of these factors contributed significantly to the model ($p > .05$) or did change the statistical result*.*

**fMRI result 1: Time-modulated neural responses over the course of aversive and appetitive acquisition**

Over the course of acquisition training, $CS_{increase\ x\ time}$ induced a greater signal increase relative to $CS_{medium\ x\ time}$ ([$CS_{increase\ x\ time} > CS_{medium\ x\ time}$]) in the primary somatosensory cortex (S1), subgenual anterior cingulate cortex (sgACC), supplementary motor area (SMA), thalamus, superior parietal lobe, frontal pole and occipital cortex, while no significant results showed neural decrease over time (Table S6).

$CS_{decrease\ x\ time}$ induced a greater signal increase over time ([$CS_{decrease\ x\ time} > CS_{medium\ x\ time}$]) in a broad range of brain regions, such as the putamen, dorsomedial prefrontal cortex (dmPFC), S1, middle frontal gyrus (MFG), precentral gyrus and occipital cortex (for details see Table S6). The search for brain regions with a decrease in activation did not yield any significant results.

**fMRI result 2: Time-modulated neural responses over the course of aversive and appetitive extinction**

During the extinction training, $CS_{increase\ x\ time}$ induced a greater signal decrease over time ([$CS_{increase\ x\ time} > CS_{medium\ x\ time}$]) in the ventromedial prefrontal cortex (vmPFC), brainstem, fusiform gyrus and occipital cortex, while no regions showed neural increase over time (Table S7).

$CS_{decrease\ x\ time}$ engaged a greater signal decrease over time ([$CS_{decrease\ x\ time} > CS_{medium\ x\ time}$]) in the vmPFC, dorsal anterior cingulate cortex (dACC), dmPFC, middle temporal gyrus and frontal pole, while the reverse contrast revealed no significant results (Table S7).

**Average neural activity underlying CS events**

***fMRI result 3: Common and distinct mechanisms of aversive and appetitive acquisition***

Regarding the average activity within acquisition training, no brain regions in the grey matter showed a comparable activation during the CS presentations ([$CS_{increase}$ - $CS_{medium}$] $\cap$ [$CS_{decrease}$ - $CS_{medium}$]).

Analysis on the differential contrast ([$CS_{increase}$ - $CS_{medium}$] > [$CS_{decrease}$ - $CS_{medium}$]) showed that, $CS_{increase}$ induced stronger activities in the brainstem, thalamus, posterior cingulate cortex (PCC) and frontal pole, and $CS_{decrease}$ yielded stronger activities in the dmPFC, dACC, precentral gyrus and S1 (Table S8).

***fMRI result 4: Common and distinct mechanisms of aversive and appetitive extinction***

In the extinction phase, no brain regions in the grey matter showed a comparable activation during presentations of the two CS types ([$CS_{increase}$ - $CS_{medium}$] $\cap$ [$CS_{decrease}$ - $CS_{medium}$]).

Compared with appetitive extinction, a stronger activation in the secondary somatosensory cortex (S2) were found during aversive extinction ([$CS_{increase}$ - $CS_{medium}$] > [$CS_{decrease}$ - $CS_{medium}$]). Appetitive extinction engaged stronger activation ([$CS_{decrease}$ - $CS_{medium}$] > [$CS_{increase}$ - $CS_{medium}$]) in areas including the MFG, parahippocampal gyrus and frontal pole (Table S8).

**Average neural activity underlying US events**

*fMRI result 5: Common and distinct mechanisms of $US_{increase}$ and $US_{decrease}$ during acquisition training*

Conjunction analysis on US events ([$US_{increase}$ $\cap$ $US_{decrease}$]) showed that a broad range of brain regions shared activation during US delivery, such as the orbitofrontal cortex, insula, dmPFC, MFG, caudate, frontal pole and occipital cortex (Table S9).

Compared with $US_{decrease}$, $US_{increase}$ engaged greater activation ([$US_{increase}$ > $US_{decrease}$]) in the areas such as the S2, dACC, thalamus, superior frontal gyrus, pallidum and occipital pole. $US_{decrease}$ elicited stronger activity ([$US_{decrease}$ > $US_{increase}$]) in areas including the S1, hippocampus, inferior parietal lobule and middle temporal gyrus (Table S10).

**Table S1. Sample characteristics**

| | Female (N=35) | Male (N=33) | Overall (N=68) |
|---|---|---|---|
| **Sex** | 35 (51.5%) | 33 (48.5%) | 68 (100%) |
| **Age** | | | |
| Mean (SD) | 37.0 (12.6) | 38.4 (13.9) | 37.6 (13.2) |
| Median [Min, Max] | 33.0 [19.0, 69.0] | 32.0 [23.0, 66.0] | 32.5 [19.0, 69.0] |
| **Temperature level for VAS40** | | | |
| Mean (SD) | 38.0 (5.19) | 39.9 (4.33) | 38.9 (4.86) |
| Median [Min, Max] | 37.5 [30.0, 45.0] | 40.5 [31.0, 45.0] | 39.9 [30.0, 45.0] |
| **Temperature level for VAS80** | | | |
| Mean (SD) | 39.3 (5.30) | 41.3 (4.48) | 40.3 (4.99) |
| Median [Min, Max] | 38.6 [30.6, 47.0] | 41.8 [32.0, 47.3] | 41.3 [30.6, 47.3] |
| **Temperature level for VAS0** | | | |
| Mean (SD) | 23.6 (6.20) | 25.8 (4.21) | 24.7 (5.40) |
| Median [Min, Max] | 22.0 [1.00, 35.0] | 26.0 [20.0, 34.0] | 24.6 [1.00, 35.0] |
| **Arousal** | | | |
| Mean (SD) | 36.7 (27.1) | 38.8 (26.4) | 37.7 (26.6) |
| Median [Min, Max] | 37.0 [0, 87.0] | 30.0 [0, 90.0] | 31.5 [0, 90.0] |
| **Pain-related fear** | | | |
| Mean (SD) | 23.2 (24.5) | 23.2 (19.9) | 23.2 (22.3) |
| Median [Min, Max] | 13.0 [0, 85.0] | 17.0 [0, 72.0] | 17.0 [0, 85.0] |
| **STADI state anxiety** | | | |
| Mean (SD) | 13.9 (3.56) | 13.6 (3.29) | 13.8 (3.42) |
| Median [Min, Max] | 14.0 [10.0, 21.0] | 13.0 [10.0, 21.0] | 13.5 [10.0, 21.0] |
| Missing | 0 (0%) | 2 (6.1%) | 2 (2.9%) |
| **STADI state depression** | | | |
| Mean (SD) | 15.6 (3.67) | 16.4 (4.67) | 16.0 (4.16) |
| Median [Min, Max] | 15.0 [10.0, 28.0] | 16.0 [10.0, 29.0] | 15.5 [10.0, 29.0] |
| Missing | 0 (0%) | 2 (6.1%) | 2 (2.9%) |
| **STADI state global** | | | |
| Mean (SD) | 29.5 (5.81) | 30.0 (6.68) | 29.8 (6.19) |
| Median [Min, Max] | 30.0 [20.0, 43.0] | 29.0 [20.0, 47.0] | 29.0 [20.0, 47.0] |
| Missing | 0 (0%) | 2 (6.1%) | 2 (2.9%) |
| **STADI trait anxiety** | | | |
| Mean (SD) | 15.6 (3.69) | 15.8 (4.44) | 15.7 (4.04) |
| Median [Min, Max] | 15.0 [10.0, 23.0] | 15.5 [10.0, 26.0] | 15.0 [10.0, 26.0] |
| Missing | 1 (2.9%) | 1 (3.0%) | 2 (2.9%) |
| **STADI trait depression** | | | |
| Mean (SD) | 14.7 (3.04) | 16.3 (3.76) | 15.5 (3.49) |
| Median [Min, Max] | 14.0 [10.0, 22.0] | 16.0 [10.0, 25.0] | 15.0 [10.0, 25.0] |

**Table S1. Sample characteristics**

| | Female (N=35) | Male (N=33) | Overall (N=68) |
|---|---|---|---|
| Missing | 1 (2.9%) | 1 (3.0%) | 2 (2.9%) |
| **STADI trait global** | | | |
| Mean (SD) | 30.3 (5.53) | 32.2 (7.11) | 31.2 (6.37) |
| Median [Min, Max] | 30.0 [21.0, 45.0] | 30.5 [20.0, 48.0] | 30.0 [20.0, 48.0] |
| Missing | 1 (2.9%) | 1 (3.0%) | 2 (2.9%) |
| **DASS anxiety** | | | |
| Mean (SD) | 0.676 (0.912) | 0.452 (0.810) | 0.569 (0.865) |
| Median [Min, Max] | 0 [0, 3.00] | 0 [0, 3.00] | 0 [0, 3.00] |
| Missing | 1 (2.9%) | 2 (6.1%) | 3 (4.4%) |
| **DASS depression** | | | |
| Mean (SD) | 1.29 (1.68) | 2.16 (1.98) | 1.71 (1.87) |
| Median [Min, Max] | 1.00 [0, 8.00] | 2.00 [0, 7.00] | 1.00 [0, 8.00] |
| Missing | 1 (2.9%) | 2 (6.1%) | 3 (4.4%) |
| **DASS stress** | | | |
| Mean (SD) | 2.62 (2.62) | 2.84 (2.96) | 2.72 (2.76) |
| Median [Min, Max] | 2.00 [0, 11.0] | 2.00 [0, 11.0] | 2.00 [0, 11.0] |
| Missing | 1 (2.9%) | 2 (6.1%) | 3 (4.4%) |
| **PSQ-20** | | | |
| Mean (SD) | 38.2 (12.9) | 40.1 (17.5) | 39.1 (15.2) |
| Median [Min, Max] | 36.7 [18.3, 66.7] | 40.0 [13.3, 80.0] | 36.7 [13.3, 80.0] |
| Missing | 1 (2.9%) | 2 (6.1%) | 3 (4.4%) |
| **PCS** | | | |
| Mean (SD) | 10.0 (9.26) | 10.9 (10.2) | 10.4 (9.67) |
| Median [Min, Max] | 8.00 [0, 34.0] | 8.00 [0, 33.0] | 8.00 [0, 34.0] |
| Missing | 1 (2.9%) | 2 (6.1%) | 3 (4.4%) |
| **PASS-20** | | | |
| Mean (SD) | 21.2 (12.4) | 27.0 (17.5) | 24.0 (15.2) |
| Median [Min, Max] | 19.5 [5.00, 55.0] | 25.0 [2.00, 71.0] | 21.0 [2.00, 71.0] |
| Missing | 1 (2.9%) | 2 (6.1%) | 3 (4.4%) |

DASS = Depression Anxiety Stress Scales; PASS = Pain Anxiety Symptom Scale; PCS = Pain Catastrophizing Scale; PSQ = Perceived Stress Questionnaire; STADI = State-Trait Anxiety-Depression Inventory; VAS = Visual Analog Scale.

**Table S2. Model description**

| Model | Included factors (FE) | Included factors (RE) | AIC | BIC | Conditional $R^2$ | Marginal $R^2$ |
|---|---|---|---|---|---|---|
| *CS valence ratings (all three conditions)* | | | | | | |
| Acquisition | CS type, time, CS type x time | Intercept: participant, Slope: CS type | 8002.3 | 8065.5 | 0.60 | 0.23 |
| Extinction | CS type, time, CS type x time | Intercept: participant, Slope: CS type | 6156.0 | 6216.6 | 0.77 | 0.24 |
| *Differential CS valence ratings* | | | | | | |
| Acquisition | CS type, time, CS type x time | Intercept: participant, Slope: CS type | 5352.8 | 5387.9 | 0.52 | 0.05 |
| Extinction | CS type, time, CS type x time | Intercept: participant, Slope: CS type | 4252.5 | 4286.2 | 0.73 | 0.02 |
| *CS-US contingency* | CS type, time, CS type x time | Intercept: participant | 4069.7 | 4101.8 | 0.30 | 0.21 |
| *US pain intensity* | US type, time, US type x time | Intercept: participant, Slope: US type | 5185.6 | 5243.0 | 0.83 | 0.68 |
| *US (un-)pleasantness* | US type, time, US type x time | Intercept: participant, Slope: US type | 5170.2 | 5227.6 | 0.80 | 0.65 |

AIC = Akaike's information criterion; BIC = Bayesian information criterion; CS = conditioned stimulus; FE = fixed effects; RE = random effects; US = unconditioned stimulus.

**Table S3. Numbers of stimuli presentations and ratings**

| Experimental phase | CS presentations | US delivery | CS valence | US pain intensity & (un)pleasantness |
|---|---|---|---|---|
| Habituation | 3x per type | - | 1x per type | - |
| Acquisition | 16x per type | 12x $US_{increase}$<br>12x $US_{decrease}$<br>24x $US_{medium}$ | 4x per type | 3x per type |
| Extinction | 12x per type | 36x $US_{medium}$ | 3x per type | 5x following $US_{medium}$<br>1x following $CS_{increase}$<br>1x following $CS_{decrease}$<br>3x following $CS_{medium}$ |

CS = conditioned stimulus; US = unconditioned stimulus.

**Table S4. Common and distinct neural responses during aversive and appetitive acquisition learning.**
Neural responses to the $CS_{increase}$ and $CS_{decrease}$ relative to the $CS_{medium}$

| | | | MNI-coordinates | | | | | |
|---|---|---|---|---|---|---|---|---|
| Contrast | Region | H | X | Y | Z | *t* | *p* | $K_E$ |
| **Common neural responses:** $[CS_{increase\ x\ time} - CS_{medium\ x\ time}] \cap [CS_{decrease\ x\ time} - CS_{medium\ x\ time}]$ | | | | | | | | |
| Neural increase over time | | | | | | | | |
| | *Occipital cortex* | *L* | *-32* | *-93* | *3* | *5.11* | *< .001* | *443* |
| Neural decrease over time | | | | | | | | |
| | - | - | - | - | - | - | - | - |
| **Distinct neural responses** (the greater, the stronger signal increase over time) | | | | | | | | |
| $[CS_{increase\ x\ time} - CS_{medium\ x\ time}] > [CS_{decrease\ x\ time} - CS_{medium\ x\ time}]$ | | | | | | | | |
| | Thalamus, MD | R | 6 | -13 | 0 | 4.06 | .042* | 2 |
| $[CS_{decrease\ x\ time} - CS_{medium\ x\ time}] > [CS_{increase\ x\ time} - CS_{medium\ x\ time}]$ | | | | | | | | |
| | - | - | - | - | - | - | - | - |

Peak voxels indicate significant activation after small volume correction using pre-defined ROIs ($p_{SVC\text{-}FWE} < .05$, ∗) or whole-brain analyses (in italic) respectively (all *p*s < .001 uncorrected). CS = conditioned stimulus; H = hemisphere; L = left; R = right; MD = mediodorsal.

**Table S5. Common and distinct neural responses during aversive and appetitive extinction learning.** Neural responses to the $CS_{increase}$ and $CS_{decrease}$ relative to the $CS_{medium}$

| Contrast | Region | H | MNI-coordinates X | Y | Z | *t* | *p* | $K_E$ |
|---|---|---|---|---|---|---|---|---|
| **Common neural responses:** $[CS_{increase\ x\ time} - CS_{medium\ x\ time}] \cap [CS_{decrease\ x\ time} - CS_{medium\ x\ time}]$ | | | | | | | | |
| Neural decrease over time | | | | | | | | |
| | vmPFC | L | -2 | 52 | -6 | 3.39 | .063 | 2 |
| Neural increase over time | | | | | | | | |
| | - | - | - | - | - | - | - | - |
| **Distinct mechanisms (the greater, the stronger signal decrease over time)** | | | | | | | | |
| $[CS_{decrease\ x\ time} - CS_{medium\ x\ time}] > [CS_{increase\ x\ time} - CS_{medium\ x\ time}]$ | | | | | | | | |
| | Parahippocampal gyrus | R | 18 | -30 | -11 | 4.02 | .041* | 2 |
| $[CS_{increase\ x\ time} - CS_{medium\ x\ time}] > [CS_{decrease\ x\ time} - CS_{medium\ x\ time}]$ | | | | | | | | |
| | - | - | - | - | - | - | - | - |

Peak voxels indicate significant activation after small volume correction using pre-defined ROIs ($p_{SVC-FWE} < .05$, ∗) (both *p*s < .001 uncorrected). CS = conditioned stimulus; H = hemisphere; L = left; R = right; vmPFC = ventromedial prefrontal cortex.

**Table S6. Time-modulated neural responses to the $CS_{increase}$ and $CS_{decrease}$ (brain regions showing a signal increase over the course of acquisition)**

| | | | MNI-coordinates | | | | | |
|---|---|---|---|---|---|---|---|---|
| Contrast | Region | H | X | Y | Z | *t* | $p_{uncorrected}$ | $K_E$ |
| [$CS_{increase\ x\ time}$ - $CS_{medium\ x\ time}$]: neural increase | | | | | | | | |
| | occipital cortex, V3 | L | -32 | -93 | 3 | 6.29 | < .001 | 808 |
| | S1 | L | -32 | -30 | 50 | 5.20 | < .001 | 100 |
| | sgACC | B | -4 | 7 | -11 | 4.86 | < .001 | 42 |
| | SMA | B | 3 | 14 | 56 | 4.66 | < .001 | 160 |
| | occipital cortex, V2 | R | 25 | -98 | 3 | 4.11 | < .001 | 128 |
| | thalamus | R | 20 | -25 | 0 | 4.09 | < .001 | 11 |
| | S1 | L | -47 | -20 | 53 | 3.84 | < .001 | 20 |
| | superior parietal lobe | L | -22 | -63 | 61 | 3.69 | < .001 | 15 |
| | frontal pole | B | -2 | 67 | 22 | 3.68 | < .001 | 16 |
| [$CS_{increase\ x\ time}$ - $CS_{medium\ x\ time}$]: neural decrease | | | | | | | | |
| | - | - | - | - | - | - | - | - |
| [$CS_{decrease\ x\ time}$ - $CS_{medium\ x\ time}$]: neural increase | | | | | | | | |
| | occipital cortex, V1 | R | 23 | -95 | 3 | 5.08 | < .001 | 85 |
| | lingual gyrus | L | -9 | -85 | -3 | 4.86 | < .001 | 603 |
| | primary motor cortex | L | -24 | -30 | 56 | 4.54 | < .001 | 141 |
| | precentral gyrus | L | -4 | -30 | 50 | 4.33 | < .001 | 18 |
| | putamen | L | -24 | -8 | 5 | 4.33 | < .001 | 18 |
| | superior parietal lobe | L | -24 | -65 | 61 | 4.33 | < .001 | 33 |
| | lateral occipital cortex | L | -22 | -83 | 17 | 4.28 | < .001 | 16 |
| | S1 | L | -12 | -45 | 67 | 4.26 | < .001 | 14 |
| | middle frontal gyrus | L | -39 | 37 | 25 | 4.26 | < .001 | 35 |
| | temporal pole | L | -57 | 9 | -9 | 4.20 | < .001 | 19 |
| | SMA | B | -2 | -11 | 50 | 4.17 | < .001 | 102 |
| | S1 | L | -49 | -25 | 45 | 4.14 | < .001 | 11 |
| | IFG | L | -54 | 14 | 14 | 4.12 | < .001 | 57 |
| | lateral occipital cortex | L | -22 | -63 | 45 | 4.10 | < .001 | 26 |
| | dmPFC | R | 6 | 17 | 50 | 4.06 | < .001 | 89 |
| | frontal pole | L | -32 | 57 | 0 | 3.79 | < .001 | 24 |
| | occipital cortex, V2 | R | 11 | -85 | -11 | 3.74 | < .001 | 13 |
| [$CS_{decrease\ x\ time}$ - $CS_{medium\ x\ time}$]: neural decrease | | | | | | | | |
| | - | - | - | - | - | - | - | - |

Peak voxels indicate significant activation thresholded at $p_{uncorrected}$ < .001 at peak-level and with a cluster size > 10 voxels. CS = conditioned stimulus; H = hemisphere; L = left; R = right; B = bilateral; S1 = primary somatosensory cortex; sgACC = subgenual anterior cingulate cortex; SMA = supplementary motor area; IFG = inferior frontal gyrus; dmPFC = dorsomedial prefrontal cortex.

**Table S7. Time-modulated neural responses to the $CS_{increase}$ and $CS_{decrease}$ (brain regions showing a signal decrease over the course of extinction)**

| | | | MNI-coordinates | | | | | |
|---|---|---|---|---|---|---|---|---|
| Contrast | Region | H | X | Y | Z | *t* | $p_{uncorrected}$ | $K_E$ |
| [$CS_{increase\ x\ time}$ - $CS_{medium\ x\ time}$]: neural decrease | | | | | | | | |
| | frontal pole/vmPFC | L | -7 | 64 | 11 | 4.30 | < .001 | 41 |
| | occipital cortex, V2 | L | -14 | -88 | -14 | 4.25 | < .001 | 22 |
| | fusiform gyrus | R | 33 | -40 | -20 | 4.13 | < .001 | 15 |
| | occipital cortex, V3 | L | -32 | -88 | -17 | 3.88 | < .001 | 14 |
| | brainstem | B | 1 | -16 | -34 | 3.84 | < .001 | 10 |
| [$CS_{increase\ x\ time}$ - $CS_{medium\ x\ time}$]: neural increase | | | | | | | | |
| | - | - | - | - | - | - | - | - |
| [$CS_{decrease\ x\ time}$ - $CS_{medium\ x\ time}$]: neural decrease | | | | | | | | |
| | middle temporal gyrus | L | -59 | -16 | -23 | 4.69 | < .001 | 74 |
| | vmPFC | B | 3 | 54 | -3 | 4.50 | < .001 | 155 |
| | dACC | R | 6 | 42 | 19 | 4.20 | < .001 | 12 |
| | dmPFC | B | -9 | 64 | 11 | 3.91 | < .001 | 59 |
| | vmPFC | B | 1 | 39 | -11 | 3.74 | < .001 | 10 |
| | frontal pole | L | -14 | 59 | 33 | 3.56 | < .001 | 13 |
| [$CS_{decrease\ x\ time}$ - $CS_{medium\ x\ time}$]: neural increase | | | | | | | | |
| | - | - | - | - | - | - | - | - |

Peak voxels indicate significant activation thresholded at $p_{uncorrected}$ < .001 at peak-level and with a cluster size >= 10 voxels. CS = conditioned stimulus; H = hemisphere; L = left; R = right; B = bilateral; vmPFC = ventromedial prefrontal cortex; dACC = dorsal anterior cingulate cortex; dmPFC = dorsomedial prefrontal cortex.

**Table S8. Distinct mechanisms of aversive and appetitive learning (average activity)**

| Contrast | Region | H | MNI-coordinates X | Y | Z | $t$ | $p_{uncorrected}$ | $K_E$ |
|---|---|---|---|---|---|---|---|---|
| Acquisition: $[CS_{increase} - CS_{medium}] > [CS_{decrease} - CS_{medium}]$ | | | | | | | | |
| | frontal pole | L | -19 | 67 | -6 | 4.01 | < .001 | 5 |
| | brainstem | B | 3 | -28 | -20 | 3.71 | < .001 | 6 |
| | thalamus | R | 13 | -8 | 3 | 3.55 | < .001 | 1 |
| | PCC | L | -4 | -38 | 19 | 3.47 | < .001 | 2 |
| Acquisition: $[CS_{decrease} - CS_{medium}] > [CS_{increase} - CS_{medium}]$ | | | | | | | | |
| | precuneus | B | 1 | -63 | 33 | 4.45 | < .001 | 53 |
| | S1 | R | 35 | -25 | 39 | 4.29 | < .001 | 8 |
| | dmPFC | R | 8 | 64 | 11 | 4.10 | < .001 | 6 |
| | precentral gyrus | L | -32 | -25 | 58 | 4.02 | < .001 | 13 |
| | S1 | R | 50 | -23 | 53 | 3.79 | < .001 | 31 |
| | precentral gyrus | R | 50 | -3 | 28 | 3.70 | < .001 | 5 |
| | dACC | R | 11 | 37 | 14 | 3.48 | < .001 | 4 |
| Extinction: $[CS_{increase} - CS_{medium}] > [CS_{decrease} - CS_{medium}]$ | | | | | | | | |
| | S2 | R | 45 | -16 | 14 | 3.24 | < .001 | 2 |
| Extinction: $[CS_{decrease} - CS_{medium}] > [CS_{increase} - CS_{medium}]$ | | | | | | | | |
| | middle frontal gyrus | L | -32 | 17 | 28 | 3.73 | < .001 | 18 |
| | frontal pole | R | 38 | 52 | -11 | 3.59 | < .001 | 4 |
| | S2 | R | 35 | -1 | 19 | 3.50 | < .001 | 8 |
| | parahippcampal gyrus | L | -17 | -35 | -14 | 3.47 | < .001 | 1 |

Peak voxels indicate significant activation thresholded at $p_{uncorrected} < .001$ at peak-level. CS = conditioned stimulus; H = hemisphere; L = left; R = right; B = bilateral; PCC = posterior cingualte cortex; S1 = primary somatosensory cortex; dmPFC = dorsomedial prefrontal cortex; dACC = dorsal anterior cingulate cortex; S2 = secondary somatosensory cortex.

**Table S9. Common neural responses to the $US_{increase}$ and $US_{decrease}$**

| Contrast | Region | H | MNI-coordinates X | Y | Z | *t* | $p_{uncorrected}$ | $K_E$ |
|---|---|---|---|---|---|---|---|---|
| $US_{increase} \cap US_{decrease}$ | | | | | | | | |
| | occipital cortex, V2 | B | 6 | -90 | 11 | 7.26 | < .001 | 1896 |
| | frontal pole | R | 48 | 44 | -6 | 5.73 | < .001 | 684 |
| | middle frontal gyrus | R | 43 | 9 | 45 | 5.41 | < .001 | 381 |
| | angular gyrus | L | -47 | -58 | 53 | 5.31 | < .001 | 368 |
| | dmPFC | B | -2 | 34 | 47 | 5.13 | < .001 | 233 |
| | inferior parietal lobule | R | 60 | -45 | 45 | 5.12 | < .001 | 501 |
| | caudate | R | 13 | 2 | 17 | 4.66 | < .001 | 112 |
| | insula | R | 40 | -11 | 5 | 4.58 | < .001 | 52 |
| | orbitofrontal cortex | L | -37 | 22 | -6 | 4.50 | < .001 | 36 |
| | caudate | L | -12 | 7 | 11 | 4.24 | < .001 | 28 |
| | middle frontal gyrus | L | -47 | 17 | 45 | 4.23 | < .001 | 109 |
| | frontal pole | L | -17 | 62 | -14 | 3.96 | < .001 | 21 |
| | insula | R | 40 | 4 | -3 | 3.94 | < .001 | 21 |
| | middle temporal gyrus | R | 53 | -25 | -9 | 3.68 | < .001 | 12 |
| | orbitofrontal cortex | R | 28 | 27 | -17 | 3.49 | < .001 | 17 |

Peak voxels indicate significant activation thresholded at $p_{uncorrected} < .001$ at peak-level and with a cluster size > 10 voxels. US = unconditioned stimulus; H = hemisphere; L = left; R = right; B = bilateral; dmPFC = dorsomedial prefrontal cortex.

**Table S10. Distinct neural responses to the $US_{increase}$ and $US_{decrease}$**

| Contrast | Region | H | MNI-coordinates X | Y | Z | *t* | $p_{uncorrected}$ | $K_E$ |
|---|---|---|---|---|---|---|---|---|
| $US_{increase}$ > $US_{decrease}$ | | | | | | | | |
| | S2 | R | 55 | 2 | 8 | 9.57 | < .001 | 1431 |
| | S2 | L | -42 | 9 | 0 | 7.14 | < .001 | 552 |
| | dACC | B | 6 | 4 | 45 | 6.82 | < .001 | 936 |
| | thalamus | B | 13 | -16 | 5 | 5.91 | < .001 | 496 |
| | precentral gyrus | R | 43 | -1 | 45 | 5.40 | < .001 | 142 |
| | occipital pole | B | -9 | -98 | -20 | 5.00 | < .001 | 278 |
| | precuneus | L | -7 | -75 | 39 | 4.54 | < .001 | 45 |
| | pallidum | L | -19 | -8 | -6 | 4.43 | < .001 | 18 |
| | frontal pole | R | 30 | 57 | 31 | 4.23 | < .001 | 29 |
| | superior frontal gyrus | L | -2 | 24 | 59 | 3.87 | < .001 | 18 |
| | S1 | R | 28 | -28 | 59 | 3.73 | < .001 | 19 |
| $US_{decrease}$ > $US_{increase}$ | | | | | | | | |
| | hippocampus | R | 25 | -11 | -20 | 6.60 | < .001 | 128 |
| | S1 | L | -49 | -28 | 64 | 6.49 | < .001 | 358 |
| | inferior parietal lobule | R | 40 | -28 | 39 | 6.07 | < .001 | 408 |
| | hippocampus | L | -24 | -16 | -20 | 5.48 | < .001 | 78 |
| | occipital cortex | R | 28 | -78 | 39 | 4.57 | < .001 | 254 |
| | middle temporal gyrus | R | 53 | -6 | -25 | 4.26 | < .001 | 27 |
| | fusiform | R | 33 | -35 | -17 | 3.97 | < .002 | 29 |

Peak voxels indicate significant activation thresholded at $p_{uncorrected} < .001$ at peak-level and with a cluster size > 10 voxels. US = unconditioned stimulus; H = hemisphere; L = left; R = right; B = bilateral; S2 = secondary somatosensory cortex; dACC = dorsal anterior cingulate cortex; S1 = primary somatosensory cortex.

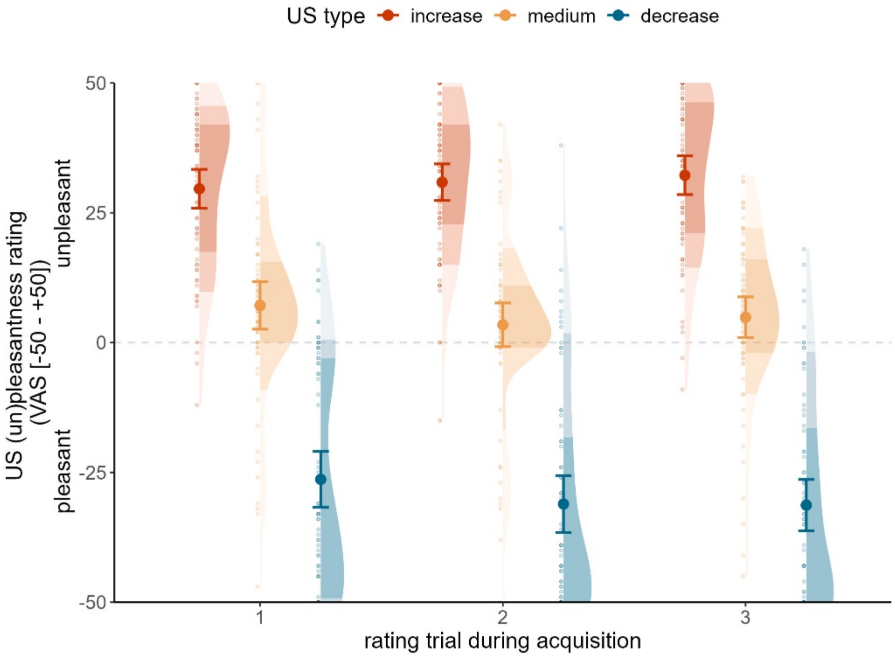


**Figure S1. US (un)pleasantness ratings.** US (un)pleasantness ratings during the acquisition training are displayed as condition-wise means (filled dots) with 95% confidence intervals, along with individual data points (open dots). Colour distribution of the violin plot represents the area into which 50 and 75% of the data points fall.
